# Strict adherence to Mendel’s First Law across a large sample of human sperm genomes

**DOI:** 10.1101/2021.11.19.469261

**Authors:** Sara A. Carioscia, Kathryn J. Weaver, Andrew N. Bortvin, Daniel Ariad, Avery Davis Bell, Rajiv C. McCoy

**Affiliations:** Department of Biology, Johns Hopkins University, Baltimore, MD, 21218, USA; School of Biological Sciences, Georgia Institute of Technology, Atlanta, GA, 30332, USA

**Keywords:** meiosis, crossovers, transmission distortion, segregation distortion, phasing, imputation

## Abstract

Mendel’s Law of Segregation states that the offspring of a diploid, heterozygous parent will inherit either allele with equal probability. While the vast majority of loci adhere to this rule, research in model and non-model organisms has uncovered numerous exceptions whereby “selfish” alleles are disproportionately transmitted to the next generation. Evidence of such “transmission distortion” (TD) in humans remains equivocal in part because scans of human pedigrees have been under-powered to detect small effects. Recently published single-cell sequencing data from individual human sperm (*n* = 41,189; 969-3,377 cells from each of 25 donors) offer an opportunity to revisit this question with unprecedented statistical power, but require new methods tailored to extremely low-coverage data (∼0.01 × per cell). To this end, we developed a method, named rhapsodi, that leverages sparse gamete genotype data to phase the diploid genomes of the donor individuals, impute missing gamete genotypes, and discover meiotic recombination breakpoints, benchmarking its performance across a wide range of study designs. After applying rhapsodi to the sperm sequencing data, we then scanned the gametes for evidence of TD. Our results exhibited close concordance with binomial expectations under balanced transmission, in contrast to tenuous signals of TD that were previously reported in pedigree-based studies. Together, our work excludes the existence of even weak TD in this sample, while offering a powerful quantitative framework for testing this and related hypotheses in other cohorts and study systems.

## Introduction

In typical diploid meiosis, each gamete randomly inherits one of two alleles from a heterozygous parent—a widely supported observation that forms the basis of Mendel’s Law of Segregation. However, many previous studies have also uncovered notable exceptions, collectively termed “transmission distortion” (TD), whereby “selfish” alleles cheat this law to increase their frequencies in the next generation. Indeed, examples of TD have been characterized in nearly all of the classic genetic model organisms, as well as numerous other systems [1–13]. Mechanisms include meiotic drive [14], gamete competition or killing [15], embryonic lethality [16], and mobile element insertion [17]. Such phenomena are frequently associated with sterility or subfertility [18,19], but may spread through the population despite negative impacts on these components of fitness [20].

Previous attempts to study TD in humans have revealed intriguing global signals but have failed to identify individual loci that achieve genome-wide significance and can be discerned from sequencing or analysis artifacts [21]. For example, using data from large human pedigrees, Zöllner et al. [22] reported a slight excess of allele sharing among siblings (50.43%)—a signature that was diffuse across the genome, with no individual locus exhibiting a strong signature. Meyer et al. [23] applied the transmission disequilibrium test (TDT) [24] to genotype data from three large datasets of human pedigrees. While multiple loci exhibited signatures suggestive of TD, the authors could not confidently exclude genotyping errors, and the signatures did not replicate in data from additional pedigrees. Similarly, Patterson et al. [25] applied the TDT to large-scale genotype data from the Framingham Heart Study but attributed the vast majority of observed signals to single nucleotide polymorphism (SNP) genotyping errors.

Genotyping of large samples of gametes offers an alternative approach for discovering TD, albeit only for TD mechanisms operating prior to the timepoint at which the gametes are collected (e.g., meiotic drive or gamete killing). Meyer et al. [23], for example, used this approach in the attempted replication of their pedigree-based test, failing to validate their most promising TD signal. Wang et al. [26] and Odenthal et al. [27] similarly used sequencing and targeted genotyping, respectively, to scan samples of sperm for evidence of TD. While neither study observed the long tracts of TD signals that are expected under a classic model of meiotic drive, they did uncover short tracts suggestive of biased gene conversion. This observation is potentially consistent with the reported rapid evolutionary turnover in the landscape of meiotic recombination hotspots [28]. Such hotspots are associated with high rates of crossovers and non-crossovers, which produce short tracts of gene conversion.

Importantly, most previous studies have been limited in their statistical power for detecting weak TD. While pedigree-based studies have been constrained by the small size of human families, gamete-based studies were historically constrained by costs and technical challenges of single-cell genotyping, limiting analysis to relatively few gametes or small portions of the genome. The recent development of large-scale whole-genome sequencing of single sperm (termed “Sperm-seq”) [29,30] offers an opportunity to study the phenomenon of TD with unprecedented statistical power. Using a highly multiplexed droplet-based approach, Sperm-seq facilitates genotyping of tens of thousands of sperm from each of several donor individuals, in turn revealing detailed patterns of meiotic recombination and aneuploidy. However, the low sequencing coverage per cell (∼0.01×) necessitates the development of tailored statistical methods for recovering gamete genotypes.

To this end, we developed a method called rhapsodi (R haploid sperm/oocyte data imputation) that uses low-coverage single-cell DNA sequencing data from large samples of gametes to reconstruct phased donor haplotypes, impute gamete genotypes, and map meiotic recombination events (Fig. 1). Here we introduce this method and quantify its performance over a broad range of sample sizes and sequencing depths, demonstrating its effectiveness across diverse study designs. Key improvements to the haplotype phasing and crossover calling methods in the Sperm-seq paper [29] include gauging model performance over a wide range of possible data inputs, directly comparing our method to an existing tool, and offering a thoroughly documented and accessible software package.

**Figure 1.**
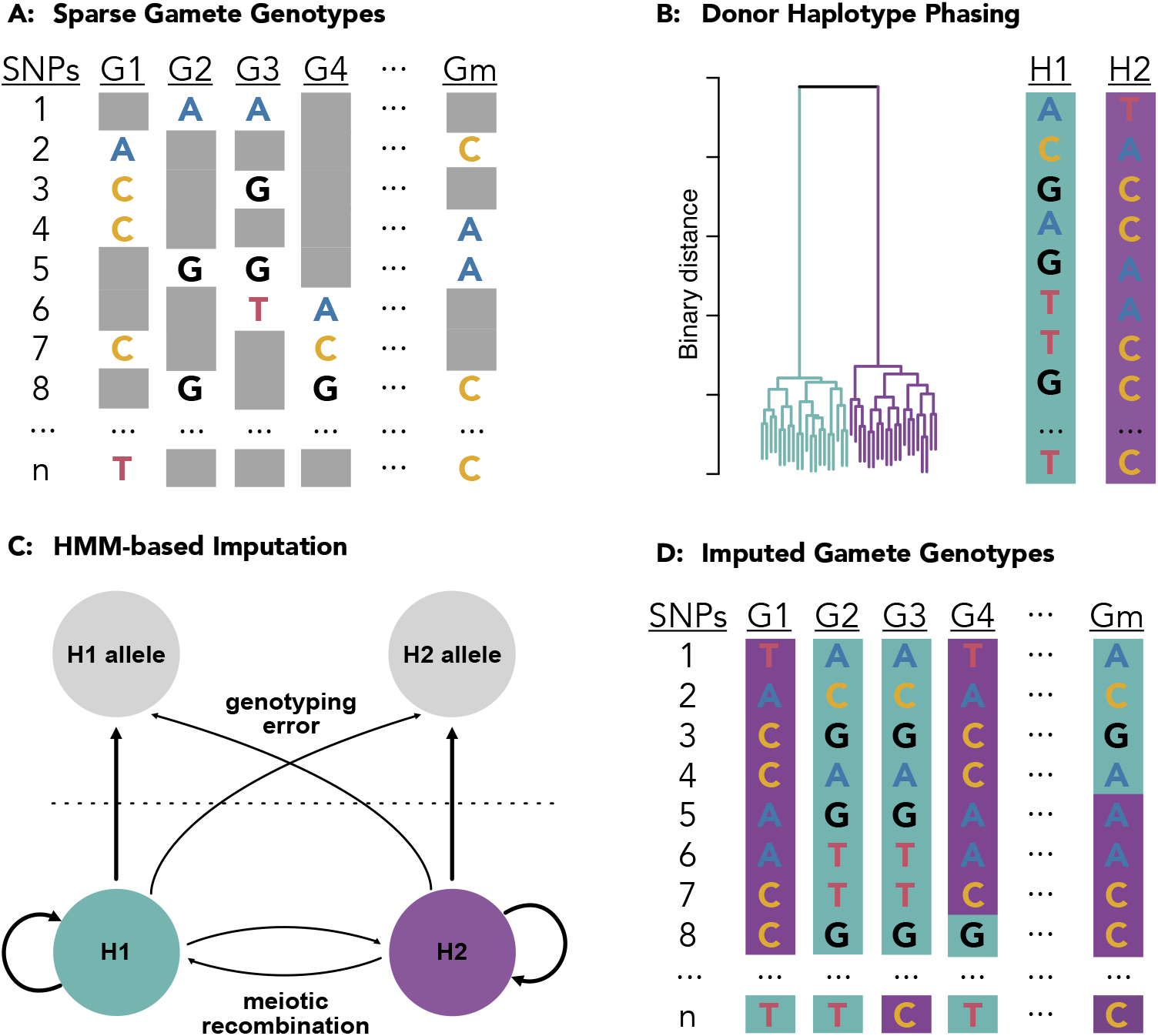
rhapsodi: R haploid sperm/oocyte data imputation. Low coverage data from individual gametes (A) is clustered to phase the diploid donor haplotypes (B). A Hidden Markov Model, with tunable rates for genotyping error and frequency of meiotic crossovers per chromosome, is applied to trace the most likely path along the phased haplotypes for each gamete (C) thereby imputing the missing gamete genotypes (D) which can be used to discover meiotic recombination events as switches from one donor haplotype to another (e.g., purple to turquoise in G4 SNPs 7 & 8 or turquoise to purple in Gm SNPs 4 & 5).

We applied rhapsodi to published single-cell sequencing data from 41,189 human sperm (969-3,377 cells from each of 25 donors) [29, 30], using the resulting imputed genotype data to scan for TD across the genome. After stringent filtering for segmental duplications and other sources of genotyping error, our results exhibited close concordance with null expectations under strict Mendelian inheritance, both with regard to individual loci and aggregate genome-wide signal. Together, our study underscores the extraordinary fidelity of human meiosis for ensuring balanced transmission of alleles to the gamete pool.

## Results

### A method for single-gamete sequencing analysis

We developed a method to phase donor haplotypes, impute gamete genotypes, and discover meiotic recombination events using low-coverage single-cell DNA sequencing data obtained from multiple gametes from a given donor (see Methods) (Fig. 1a). Briefly, chromosomes are segmented into overlapping windows, and within each window, the sparse gamete genotype observations at heterozygous SNPs (hetSNPs) are clustered to distinguish the two haplotypes (i.e., phase the genotypes) of the diploid donor individual (Fig. 1b). The sequences of alleles that compose these haplotypes are decoded based on majority votes within each cluster, and haplotypes from overlapping windows are then stitched together based on sequence identity, thereby achieving chromosome-scale phasing. A Hidden Markov Model (HMM), with transition probabilities determined by rates of meiotic crossover and emission probabilities determined by rates of genotyping error, is then used to infer the most likely path along the phased haplotypes for each gamete (Fig. 1c), thereby imputing missing genotype data (Fig. 1d). Points where such paths are inferred to switch from one donor haplotype to the other suggest the locations of meiotic recombination events.

### Evaluating performance on simulated data

In order to benchmark rhapsodi’s performance, we developed a generative model to construct input data with varying sample sizes, rates of meiotic recombination, sequencing coverages, and genotyping error rates (Fig. S1). We then applied rhapsodi to the simulated data, matching input parameters to those used in the simulations (average of 1 recombination event per chromosome and genotyping error rate of 0.005). Phasing was assessed based on accuracy, completeness, switch error rate, and largest haplotype segment (Fig. 2a, Fig. S2a). Across the range of study designs, we observed that phasing performance improved with increasing amounts of data (gamete sample size and coverage). For specific scenarios involving low coverages or small numbers of gametes, this relationship was not always monotonic. This suggests that other parameters that we currently hold fixed (e.g., window size used in phasing) may interact and influence performance—an area for future methodological fine-tuning.

**Figure 2.**
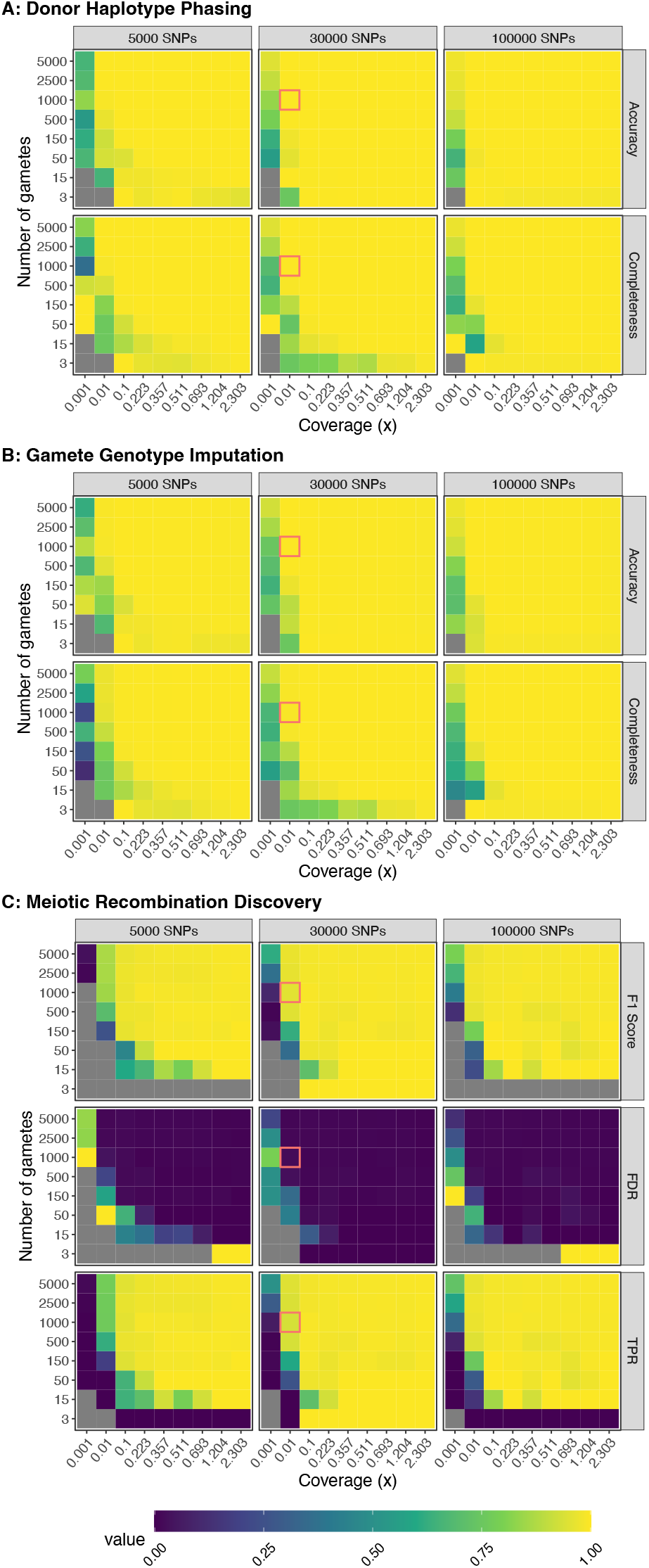
Benchmarking performance across a range of study designs. Values represent the average of three independent trials. FDR: False Discovery Rate; TPR: True Positive Rate. For phasing and imputation, gray indicates that no hetSNPs remained after downsampling. For discovery, gray indicates the absence of a prediction class (e.g., zero FNs, FPs, TNs, or TPs). Simulations roughly matching the characteristics of the Sperm-seq data are outlined in red.

With the exception of very small gamete counts, we observed that phasing performance reaches a plateau at ∼0.1 × coverage. Yet performance is also high at even lower coverages when compensated by large sample sizes of gametes. Qualitatively similar trends were observed for the tasks of imputing gamete genotypes and discovering meiotic recombination breakpoints (Fig. 2b and Fig. 2c; Fig. S2b and Fig. S2c).

For study designs roughly mirroring the published Sperm-seq data [29] (1,000 gametes, 30,000 hetSNPs, coverage of 0.01×), rhapsodi phased the donor haplotypes with 99.993 (±0.003)% accuracy and 99.96 (±0.03)% completeness (Fig. 2a); imputed gamete genotypes with 99.962 (±0.002)% accuracy and 99.34 (±0.04)% completeness (Fig. 2b); and discovered meiotic recombination breakpoints with a mean F1 Score of 0.959 (±0.003), a mean False Discovery Rate (FDR) of 1.8 (± 0.3)%, and a mean True Positive Rate (TPR) of 93.7 (±0.3)% (Fig. 2c and Fig. S2c) (values reported as the mean, plus or minus one standard deviation; see Supplemental Text for definitions of metrics).

Discovery of meiotic recombination events exhibited the weakest relative performance among the three tasks, albeit still strong in absolute terms. This is likely due to a combination of this task’s dependence on the successful completion of the previous two tasks, the simplifying assumptions employed within the generative model, and the inherently challenging nature of this task in data-limited scenarios. In relation to the last point, we observed that the resolution of inferred meiotic recombination breakpoints was strongly associated with the sequencing coverage of the input data (Fig. S3a), as well as the underlying density of hetSNPs across the genome. Assuming a pairwise nucleotide diversity of 1 per 1,000 bp [31], and given that the theoretical limit of median resolution is two hetSNPs, we found that a coverage of 2.3× (missing genotype rate of 10%) was required to approach this theoretical limit. Meanwhile, for coverage in line with the Sperm-seq data [29] (∼0.01×), we inferred a median resolution of 167.5 kbp, in line with empirical observations from the original study [29].

Through closer investigation of the locations of false positive (FP) and false negative (FN) recombination events (Fig. S3b), we identified three error modes that originate from various limitations of the data. Specifically, we attributed the vast majority of FNs to crossovers occurring near the ends of chromosomes or pairs of crossovers occurring in close proximity to one another, especially at low coverages (Fig. S3c). In the second of such cases (co-occurring FNs), the genotype data may be too sparse to capture one or more informative markers that flank the recombination breakpoint(s). Notably, such nearby crossovers should be rare in data from humans and many other species due to the phenomenon of crossover interference [32], which causes crossovers to be spaced farther apart than expected by chance. By simulating crossover locations under a uniform distribution, our benchmarking strategy is thus conservative, as it would overweight the importance of this error mode. A third mode of error, which manifests as pairs of FPs and FNs, owes to slight displacement of the crossover breakpoint (Fig. S3c), which may arise as a consequence of premature or delayed switching behaviors of the HMM.

We next assessed rhapsodi’s robustness to parameter mis-specification by altering the recombination and genotyping error rate parameters relative to the simulations. Only one parameter was mis-specified at a time, and the other was matched to that used in the simulation. While our results suggest overall robustness, we found that overestimating the genotyping error rate or average recombination rate (Fig. S4 and Fig. S5) had less effect on performance than underestimating either of these parameters (Fig. S6 and Fig. S7). In practice, such parameters may be informed based on outside knowledge for a given species (recombination rate ≈ 1 × 10^−8^ per bp for humans) and sequencing platform (error rate ≈ 0.005 per bp).

### Benchmarking against existing methods

We next compared rhapsodi to an existing software tool, called Hapi, which was recently developed for the same task of gamete imputation and was demonstrated to outperform all similar methods [33]. We compared the performance of rhapsodi and Hapi using simulated data from a range of 3 to 150 gametes and coverages ranging from 0.001× to 2.303× (Fig. 3). Datasets with more than 150 gametes were not benchmarked because Hapi’s runtime with larger sample sizes became intractable, taking up to 39 hours per simulated dataset. Of 2,916 simulated datasets, Hapi was able to successfully phase, impute, and detect crossovers in 1,902 datasets (65%), while rhapsodi was able to do so in 2,754 datasets (94%) (Fig. S8). For datasets that Hapi could not analyze, rhapsodi maintained high accuracy and completeness (Fig. S9) with low cost to performance in comparison to datasets that both tools analyzed successfully. Hapi typically achieved high phasing accuracy, but often at the cost of low completeness (Fig. 3a). In contrast, rhapsodi exhibited relative balance between accuracy and completeness. Across a large range of study designs, rhapsodi phased and imputed a greater proportion of SNP genotypes than Hapi with little cost to accuracy (Fig. 3a – Fig. 3b). This increase in completeness was most pronounced at the low coverages (<0.1×) that characterize the Sperm-seq data (Fig. 3).

**Figure 3.**
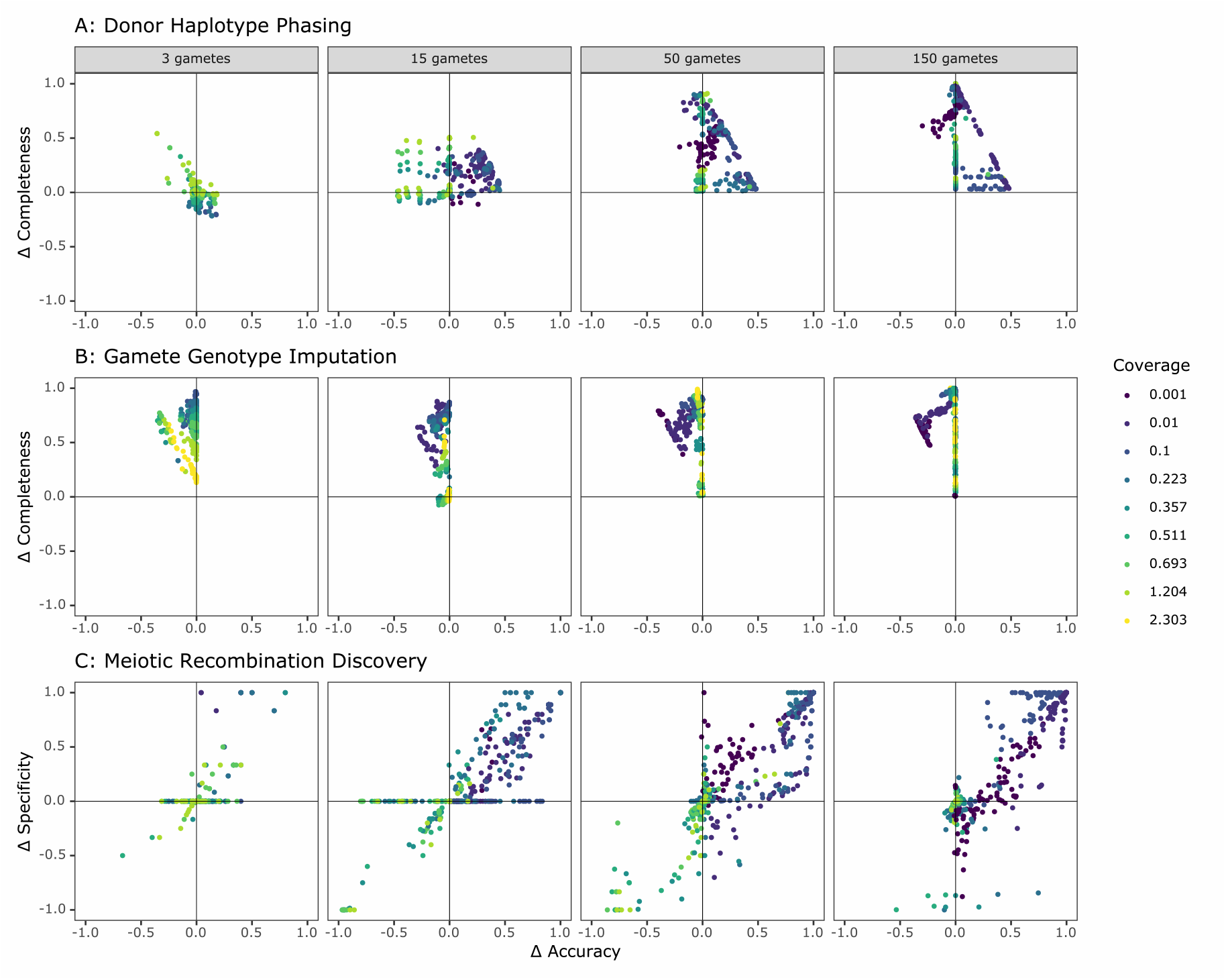
Comparison of rhapsodi performance with Hapi, an existing gamete imputation tool. Figure shows difference in performance between both tools (rhapsodi - Hapi) on simulated data. Each point represents a simulated dataset, and only datasets successfully analyzed by both tools are displayed.

### Application to data from human sperm

Given this confidence in the performance of our method, we proceeded to the analysis of published [29,30] single-cell DNA sequencing data from 41,189 human sperm (969-3,377 cells from each of 25 donors; ∼0.01× coverage per cell) (Fig. S10). Before applying rhapsodi, we performed stringent filtering to mitigate sequencing, alignment, and genotyping errors that could confound our downstream tests of TD. Specifically, we removed low-quality and aneuploid cells for the chromosome of interest; excluded regions of low mappability or extreme repeat content; restricted analysis to known SNPs from the 1000 Genomes Project [34]; and excluded regions harboring potential segmental duplications in each donor individual’s genome (see Methods and Supplemental Text). While this stringent filtering removed a total of 15,138,461 (∼30%) SNPs from the input data, we emphasize that true signatures of TD should be minimally affected by such filtering, as such signatures are expected to extend across large genomic intervals due to the extreme nature of linkage disequilibrium (LD) among the sample of gametes (Fig. S11).

Applying rhapsodi to these filtered data, we phased donor haplotypes at an average of 99.9 (±0.16)% of SNP positions per donor and chromosome (i.e., leaving ∼0.1% as unknown phase); imputed an average 99.3 (±1.23)% of genotypes per gamete and chromosome; and identified an average of 1.17 (median of 1) meiotic recombination events per gamete and chromosome, with a mode of 24 autosomal crossovers per gamete genome, broadly consistent with the reported length of the autosomal genetic map for males [35]. The average resolution of breakpoints was 664 kbp (±1.25 Mbp), with a median of 357 kbp. (Values are reported as the mean, plus or minus one standard deviation.) This median is in line with empirical observations from the original study [29]. As previously reported [29], the inferred crossover locations were concentrated in similar genomic regions across all 25 donors, with the highest densities occurring near the ends of chromosomes (Fig. S12). The overall distributions of crossover locations were qualitatively similar to the patterns observed in the prior analysis of a subset of these donors [29].

### Strict adherence to Mendelian expectations across sperm genomes

The scale of the Sperm-seq dataset offers strong statistical power to detect even slight deviations from Mendelian expectations, as supported by our simulations of TD across a range of gamete sample sizes and transmission rates (Fig. S13). For each of the 25 donors, we scanned across the imputed sperm genotypes, performing a binomial test for each SNP. Naive multiple testing approaches based on the nominal number of tests would be over-conservative in this context, given the extreme levels of LD across the sample of closely related sperm. We therefore applied a principal component analysis-based approach to infer the effective number of independent statistical tests [36, 37] and used this value as the basis of a Bonferroni correction (p-value threshold = 1.78 × 10^−7^; see Methods). After applying this correction, no individual SNP exhibited evidence of TD at the level of genome-wide significance (Fig. 4).

**Figure 4.**
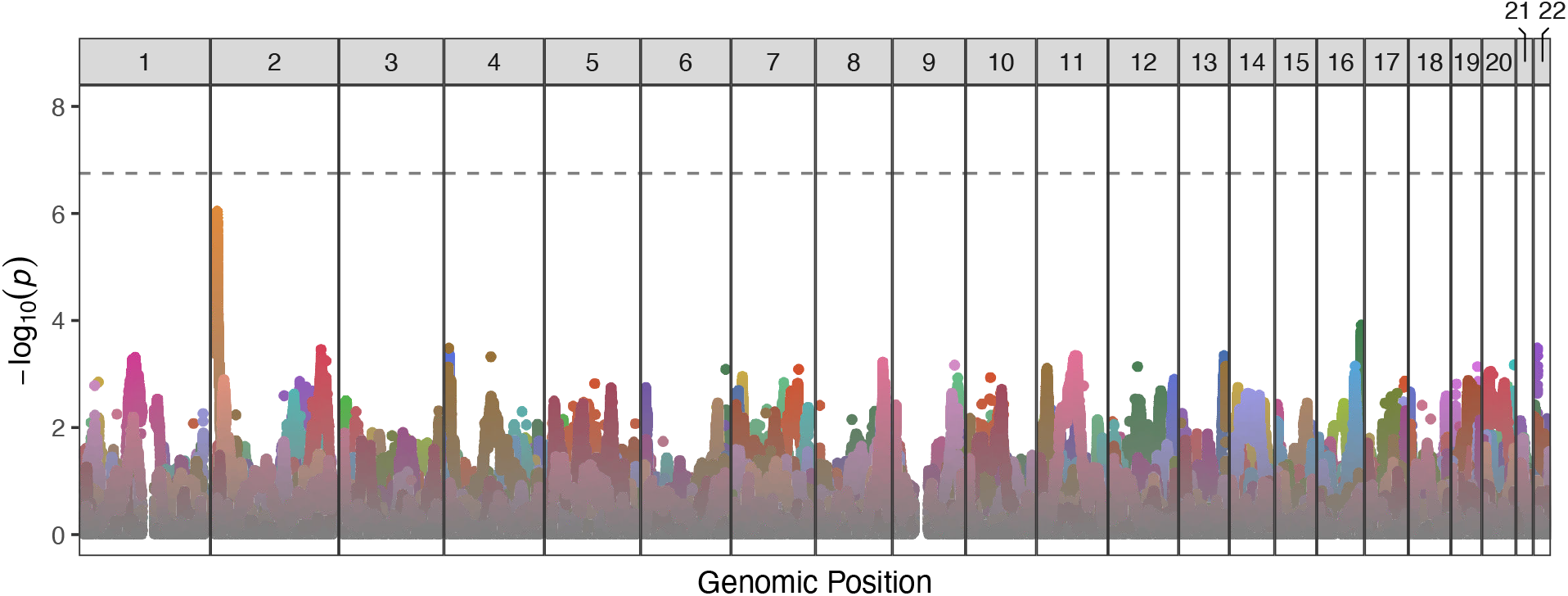
Manhattan plot displaying genome-wide TD scan results for all 25 sperm donors across the 22 autosomes. P-values are correlated across large genomic intervals due to the high degree of linkage disequilibrium among sperm cells from a single donor. Colors distinguish results from different donors. No individual test was significant after multiple testing correction (p-value threshold = 1.78 × 10^−7^).

The strongest signal of TD occurred at the end of chromosome 2 (transmission rate = 833 of 1571 [56.2%]; p-value = 9.6 × 10^−7^) for donor NC18 (depicted in dark yellow in Fig. 4) and encompassed 7 SNPs (rs7603674, rs7578293, rs7567762, rs12478306, rs2084684, rs2385305, rs2100268) within this gene poor region. These linked SNPs segregate at frequencies of ∼0.25 in European populations of the 1000 Genomes Project (of greatest relevance given the reported European ancestry of that donor) [34, 38].

### No global signal of biased transmission in human sperm

In addition to the absence of locus-specific signatures achieving genome-wide significance, we observed that the combined distribution of p-values closely mirrors that which is expected under the null hypothesis of no TD (Fig. 5). This absence of global signal is intriguing given the strong statistical power underlying the analysis.

**Figure 5.**
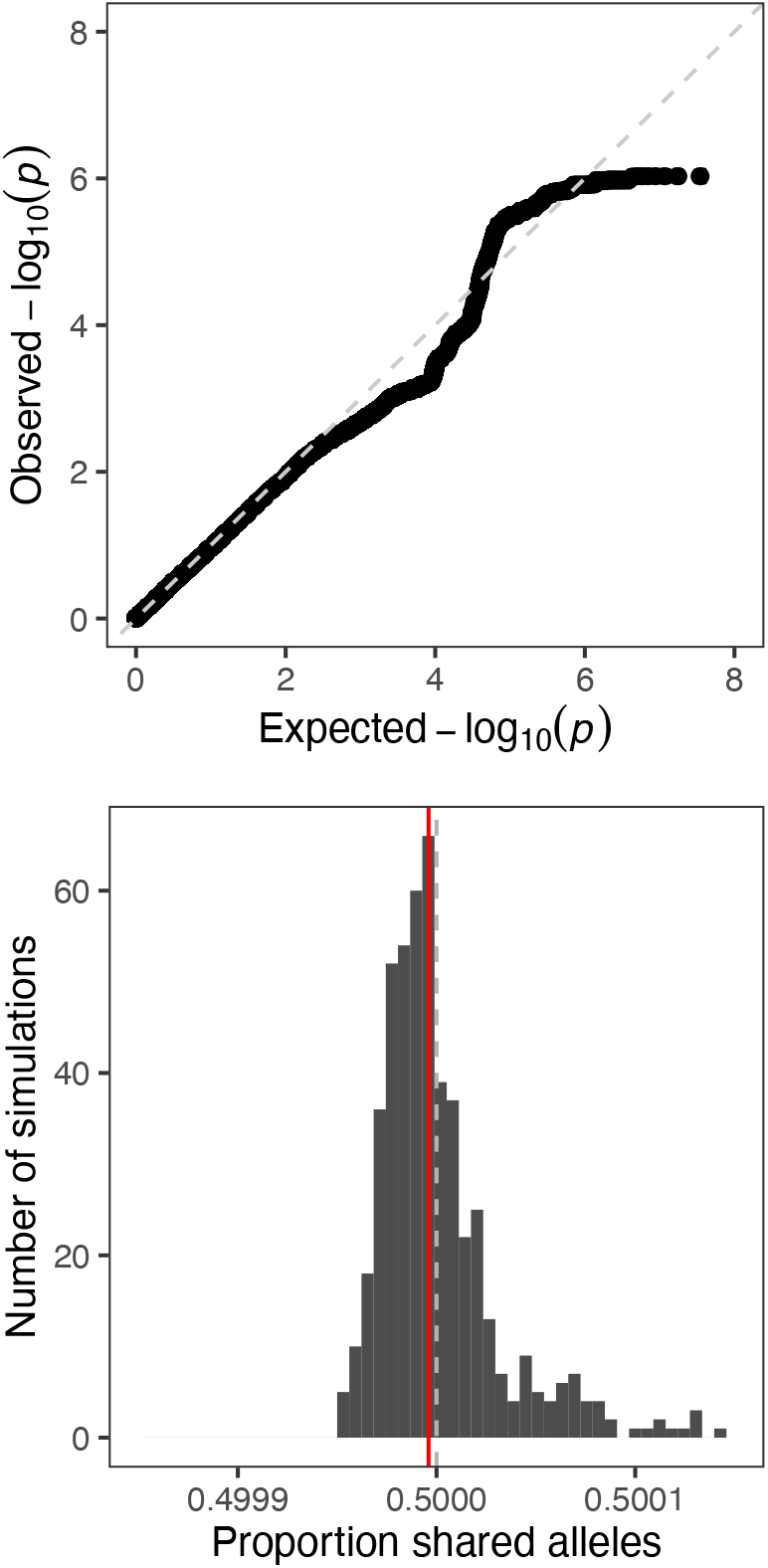
Quantifying global evidence of TD. (A) Quantile-quantile plot comparing the distribution of p-values from our genome-wide scan for TD to that expected under the null. The dashed line corresponds to x = y. (B) Mean allele sharing across all pairs of sperm from null simulations (n = 500) is depicted with grey bars. The same mean computed on the observed data (0.499996) is depicted with a red line, while the null proportion of 0.5 is depicted with a gray dashed line.

To more formally quantify global evidence of TD, we calculated the overall degree of allele sharing at hetSNPs among sperm from the same donor. After pruning the data to SNPs in approximate linkage equilibrium (see Methods) we found that across all pairwise comparisons of sperm from each of 25 donors, the average proportion of shared alleles was 0.499996, consistent with the distribution of this proportion under null simulations with no TD (n = 500 simulations; p-value = 0.47; Fig. 5).

## Discussion

Past studies of TD in humans have been limited in statistical power, either due to the small size of human families or costs and technical limitations of gamete genotyping. Such limitations are compounded by the high burden of multiple testing in genome-wide scans. Here we developed a genotype phasing and imputation method, termed rhapsodi, that allowed us to leverage low-coverage sequencing data from a large number of single gametes to impute gamete genotypes and test for TD with unprecedented statistical power.

In contrast to the tentative locus-specific and genome-wide signals reported in previous studies, our results are consistent with strict adherence to Mendel’s Law of Segregation across this sample of human sperm genomes. The signals that we observed are well explained based on the expected variance of the binomial process of Mendelian segregation under the null hypothesis of balanced transmission genome-wide. This negative result hinged on stringent filtering of the input data, as phenomena such as hidden structural variation and ensuing mapping and genotyping errors may otherwise generate false signatures of TD. It is important to emphasize that even stringent filtering is not expected to eliminate true signals, which should span large genomic intervals due to the extreme LD within the samples of sperm from a given donor.

Given the strong statistical power of our study, the absence of TD signal excludes the possibility of even weak TD. This negative result provides a striking contrast to the numerous validated examples uncovered in other species. However, our observations do not preclude the occurrence of TD in humans, for reasons that we enumerate below. First, our study was limited to sperm from 25 donor individuals. If true TD involves rare alleles or particular patient populations that were not sampled in the datasets we analyzed, our study design would miss such cases. A second and related consideration is that if not balanced by countervailing forces, alleles that are subject to TD may rapidly fix in the population, such that detecting such alleles during their ephemeral existence as polymorphisms may be exceedingly unlikely. Third, our analysis is solely sensitive to mechanisms of TD that would operate prior to the sampling of the sperm cells (e.g., sperm killing), and is therefore blind to mechanisms that may operate in female meiosis (e.g., meiotic drive) or at different stages of development (e.g., fertilization, maternal-fetal incompatibility, postzygotic selection, etc.). Notably, female meiosis may be particularly susceptible to meiotic drive given the asymmetric nature of the cell divisions that produce the oocyte and polar bodies [39]. Finally, our analysis does not exclude the existence of TD driven by biased gene conversion, which would generate short tracts below the resolution of our study. Indeed, the occurrence of this phenomenon is well documented based on both comparative evolutionary data and targeted analysis of human recombination hotspots, and is beyond the scope of our study [28].

## Conclusion

In summary, we introduced and benchmarked a method to phase donor haplotypes, impute gamete genotypes, and infer meiotic recombination events using low-coverage single-cell sequencing data from a large sample of gametes. We applied this method to scan a large sample of human sperm genomes for evidence of “selfish” alleles that violate Mendel’s Law of Segregation. We observed no evidence of even weak effects across this sample, either with respect to individual loci or aggregate genome-wide signal, in turn supporting a strict model of balanced transmission. Measuring the fidelity of Mendelian segregation and uncovering the mechanisms that safeguard or subvert this process remain important goals for human genetics. Given the documented fertility impacts of TD in other organisms, a thorough understanding of the loci driving such inheritance patterns could help reveal novel genetic causes of human infertility. More broadly, our method offers a flexible and reproducible toolkit for testing this and related hypotheses in other cohorts and study systems, especially in light of the declining costs and expanding application of single-cell sequencing methods.

## Methods

### Data input and filtering

The main algorithmic steps of rhapsodi consist of (1) data preprocessing and filtering, (2) reconstruction of phased donor haplotypes (Fig. 1b), (3) imputation of gamete genotypes (Fig. 1c), and (4) discovery of meiotic recombination breakpoints. A table of sparse gamete genotypes (SNPs × gametes) with corresponding genome positions is required by rhapsodi as input (Fig. 1a). Genotypes may be encoded in one of two forms: 0/1/NA (denoting reference, alternative, or missing genotypes, respectively) or A/C/G/T/NA (denoting nucleotides or missing genotypes, respectively). For A/C/G/T/NA encoding, reference and alternative alleles are also required as input. Data is then filtered to consider only SNP positions that are heterozygous for the given individual, with at least one reference and one alternative allele observed across gametes. Phased donor haplotypes, imputed genotype (and corresponding source haplotype) for each SNP in each gamete, and a list of locations of recombination breakpoints detected for each gamete is returned by rhapsodi as output (Fig. 1d). We detail the methods that produce each output below.

### Reconstruction of phased donor haplotypes

Our phasing approach leverages patterns of co-inheritance across gametes to reconstruct donor haplotypes from the sparse genotype matrix derived from low coverage sequencing (Fig. 1a). Chromosomes are divided into overlapping windows, and within each window, we compute pairwise binary (Jaccard) distances, cluster the gamete genotypes using Ward’s method [40] (implemented with the “ward.D2” function in R), and cut the resulting tree into two groups to distinguish the haplotypes (Fig. 1b). For each cluster within a window, we use majority voting to decode the phased haplotype sequences. This approach depends on the reasonable assumption that recombination is suffciently rare, affecting a small minority of gametes within each window. If no genotype achieves a majority at a given position, we designate the phase at that position as ambiguous (denoted as “NA”). Finally, haplotypes of adjacent (overlapping) windows are stitched together by considering the genotype consensus (i.e., rate of matching) for the overlapping SNPs. In the stringent stitching mode, if consensus is greater than a given threshold (default = 0.9), the haplotypes under consideration are inferred to be the same (i.e., to derive from the same donor homolog). If the consensus is less than 1 minus the threshold (default = 1 -0.9 = 0.1), the haplotypes under consideration are inferred to be distinct (i.e., to derive from different donor homologs). Meanwhile, if the consensus lies between these upper and lower boundaries, the windows are considered too discordant to ascertain the relation of the haplotypes, and the method returns an error. However, rhapsodi can optionally be run in a lenient stitching mode which continues with phasing regardless of the level of consensus. In this case, the haplotypes with the highest consensus are assumed to derive from the same homolog.

### Imputation of gamete genotypes

Our imputation approach uses a Hidden Markov Model (HMM) and a subsequent filling step to infer the genotypes of each gamete, effectively tracing the most likely path through the donor haplotypes given the sparse observations for a given gamete. Our model assumes that (1) the hidden haplotype state at a given SNP depends only on the haplotype state at the SNP immediately prior; (2) adjacent genotypes originate from the same donor haplotype unless a meiotic recombination event occurred between the two; (3) the observed SNP alleles are either correct or in rare cases arise from genotyping errors (e.g., PCR errors, contaminant DNA co-encapsulated with the sperm genome during library preparation, sequencing errors, mapping errors, etc.). The HMM uses the phased donor haplotypes and accounts for probabilities of genotyping errors and meiotic crossovers: the probability of genotyping errors is reflected in the emission probabilities, while the probability of meiotic crossovers is reflected in the transition probabilities (Fig. 1c). We then apply the Viterbi algorithm [41] to determine the most likely sequence of hidden states, iterating over all gametes independently (Fig. 1d). To avoid potential over-smoothing behavior of the HMM, we optionally overwrite the imputed alleles with any observed alleles for rare sites where these genotypes conflict. The HMM only considers observed genotypes and makes no predictions for missing genotypes. To impute missing genotypes, we assume that tracts of unobserved sites that are bordered by SNPs originating from a single donor haplotype also originate from that same haplotype. We thus fill in the genotypes for these unobserved sites. Similarly, we assume that recombination does not occur between the last observed SNPs and the ends of the chromosomes, again filling genotypes at unobserved sites in these terminal regions. Tracts of unobserved sites that are bordered by SNPs assigned to different donor haplotypes remain unassigned (denoted as “NA”).

### Discovery of meiotic recombination breakpoints

Using the imputed gamete genotypes and the underlying phased donor haplotypes, we identified meiotic recombination breakpoints as sites where, for a given gamete chromosome, the inferred path switches from one donor haplotype to the other. This step should typically be run on the smoothed data (i.e., without superimposing the original observations), as even a low rate of genotyping error could otherwise manifest as false meiotic recombination. Recombination breakpoints are reported as intervals, defined by the last observed site inferred to derive from one haplotype and the first observed site inferred to derive from the other. These start and end points thus demarcate regions in which the meiotic recombination events likely occurred.

### Assessing performance with simulation

We developed a generative model to simulate input data and assess rhapsodi’s performance (Fig. S1) while varying (a) the number of gametes, (b) the underlying number of hetSNPs, (c) depth of coverage, (d) recombination rate, and (e) genotyping error rate. Given these arguments, we construct a sparse matrix for input to the rhapsodi pipeline. The algorithmic steps of this generative model include (1) building phased diploid donor haplotypes (Fig. S1a), (2) randomly tracing through these haplotypes in the face of recombination to construct gamete genotypes (Fig. S1b), (3) masking genotype observations to mimic low sequencing coverage (Fig. S1c), (4) superimposing genotype errors at a given rate (Fig. S1d), and (5) filtering to only retain hetSNPs. We detail each step in turn below.

In the first step of the generative model, we construct phased haplotypes of the diploid donor by randomly generating a vector of 0’s and 1’s (with equal probabilities) at a length equal to the underlying number of hetSNPs. This binary vector is then inverted to generate the complementary haplotype (Fig. S1a).

We then use these simulated homologs to generate genotypes for each gamete (Fig. S1b). Specifically, we sample a count of recombination breakpoints from a Poisson distribution, where lambda reflects the average recombination rate. Each recombination breakpoint is randomly placed on the respective chromosome, and genotypes occurring after that breakpoint are inverted. In cases where a gamete chromosome does not recombine, the chromosome is identical to one of the parental chromosomes. By placing crossovers randomly, our method ignores heterogeneity in the landscape of meiotic recombination, as well as the phenomenon of crossover interference [32], which suppresses nearby crossovers.

After obtaining the gamete genotype matrix, we simulated the sparse coverage of single-cell DNA-seq by masking observed genotypes. The missing genotype rate (MGR) is obtained from the probability density function of a Poisson distribution with x = 0 and lambda equal to the coverage metric. We then multiplied the MGR by the total number of simulated SNP observations (number of underlying hetSNPs × number of gametes) to compute the total number of genotypes that should be masked (denoted as “NA”) (Fig. S1c). Following a similar approach, we computed the number of genotyping errors by multiplying the error rate by the number of nonmissing genotypes and randomly placed these errors at individual SNPs by inverting the respective alleles (0 to 1 or vice versa) (Fig. S1d).

Finally, SNPs or rows with at least one 0 genotype and one 1 genotype are retained, and the rest of the rows are filtered out, retaining only hetSNPs where both alleles were observed. In this way, the final number of hetSNPs in the generated output may be less than or equal to the specified input number of underlying hetSNPs.

We used this generative model to construct input data for ranges of gamete sample sizes (3, 15, 50, 150, 500, 1000, 2500, 5000), underlying SNPs (5000, 30000, 100000), and depths of coverage (0.001, 0.01, 0.1, 0.223, 0.357, 0.511, 0.693, 1.204, and 2.303×, corresponding to missing genotype rates of 99.9, 99, 90, 80, 70, 60, 50, 30, 10%, respectively). For each combination of these three parameters, nine different conditions were simulated, with three independent runs for each condition. The conditions were defined by combinations of the expected genotyping error (0.001, 0.005, and 0.05) and average recombination rate (0.6, 1, and 3) used to construct the gamete data. Simulations in which 0 hetSNPs remained after filtering were dropped from downstream benchmarking.

We evaluated the quality of donor haplotype phasing by considering switch error rate, largest haplotype segment, completeness, and phasing accuracy (see Supplemental Text for definition of metrics). To assess the quality of imputation of gamete genotypes, we similarly quantified accuracy, completeness, and switch error rate. Finally, we assessed meiotic recombination discovery by computing the True Positive Rate (TPR), the False Discovery Rate (FDR), and the breakpoint resolution. Given the imbalance in class membership for the number of true recombination breakpoints compared to the number of false recombination breakpoints, we also considered precision and F1 Score.

### Comparison to existing methods

We compared our method to the software package Hapi (haplotyping with imperfect genotype data) [33], which was similarly developed for phasing and imputation based on single-cell DNA sequencing of gametes. Hapi was demonstrated to outperform the only other haploid-based algorithm, PHMM (pairwise Hidden Markov Model) [42], as well as two commonly applied diploid-based phasing methods, WhatsHap [43] and HapCUT2 [44] in terms of accuracy, reliability, completeness, and cost-effectiveness. Because Hapi outperforms PHMM and the PHMM software was not readily available, we did not directly compare PHMM with rhapsodi.

In contrast to our method, Hapi was designed for datasets that contain relatively small numbers of gametes (∼3-7) sequenced at individually higher coverages (> 1×). Given this distinction, the default behavior of Hapi’s autorun function was not appropriate for our simulated datasets. In particular, Hapi’s imputation function (hapiImpute) by default filtered out any SNPs that had a missing genotype observation for at least one gamete, which results in all data being removed from our low-coverage datasets. We thus disabled this filtering behavior prior to benchmarking against our method.

As part of Hapi’s autorun function, the HMM used to filter errors uses a fixed set of transmission and emission probabilities. To more fairly compare the tools to each other, we instead provided Hapi with an HMM with parameters matching those of our simulation, thus closely mirroring the approach we used to evaluate rhapsodi in this analysis. After rhapsodi and Hapi were run on a dataset, the performances of both methods were evaluated using rhapsodi’s assessment functions.

### Genome-wide scan for TD

To scan the genome for TD, single-cell genotype calls were generated from sperm sequencing data as described by Bell et al. [29]. Briefly, genomic variants were called for each donor using all sequence data from all sperm cells. After selection of heterozygous sites, the software GenotypeSperm (Drop-seq Tools v2.2, https://github.com/broadinstitute/Drop-seq/releases) was used to determine which sites were captured by each sperm cell barcode and which allele(s) were present at each of these observed sites. Aneuploidy status and cell doublets were identified as previously described [29,30]. Only cell barcodes associated with single sperm cells (non-doublets) with even sequence coverage were included in the analysis, while aneuploid autosomes were excluded from the analysis.

Before applying rhapsodi for phasing and imputation, we applied several additional filtering criteria with the goal of restricting analysis to genuine SNPs and mitigating artifacts induced by sequencing, mapping, and other sources of genotyping error. Specifically, we excluded all technically challenging regions of the genome previously identified by the Genome in a Bottle Consortium (GRCh38 “union”) [45] and/or the ENCODE Consortium [46]. We additionally required at least one observation of both the reference and alternative allele across sperm from a given donor. We also restricted to SNPs that were previously observed to be polymorphic in external data from the 1000 Genomes Project [34]. Finally, we removed any sites with evidence of segmental duplication (see Supplemental Text and Fig. S14).

We used a two-tailed binomial test (comparing counts of each allele under a null hypothesis of balanced transmission) to scan for TD at each hetSNP across sperm genomes from each of 25 donors. As input to this analysis, we used the imputed gamete data output by rhapsodi, applying the option to superimpose any observed genotypes (at rare sites of discordance with inferred HMM state).

### Significance threshold for TD scan

In our previous step, we applied a separate binomial test for TD at each hetSNP in each donor individual. Given the relative infrequency of recombination (per base pair per meiosis), sperm genomes are composed of large tracts of the donor’s two parental haplotypes. By consequence, genotype data from the sample of sperm from a given donor are affected by extreme LD, which typically extends across entire chromosomes. As such, our binomial tests are highly correlated across the genome, and naive multiple testing correction based on the number of SNPs would be overly conservative.

To account for this effect, we applied the method simpleM [36,37], which uses a principal components analysis (PCA) approach to compute the effective number of independent statistical tests. For the Sperm-seq dataset, simpleM estimated 281,368 effective tests (compared to 34,799,282 nominal tests). We then used the effective number of independent tests to apply a Bonferroni correction of 0.05 / 281,368 = 1.78 × 10^−7^.

### Calculation of global signal of TD

We quantified global evidence of TD across the sperm genomes by calculating the rate of allele sharing all pairs of sperm from each donor. First, we applied PLINK to the fully imputed sperm genotype matrix output by rhapsodi to generate a list of the SNPs deemed to segregate in near linkage equilibrium (plink –file filename –no-fid –no-parents –no-sex –no-pheno –indep 50 5 2 –out out-filename) [47]. We then computed a Hamming distance matrix, effectively counting mismatches between each pair of sperm genomes from each donor. After repeating the above steps for all donors, we reported the mean across all pairs of sperm cells.

Null simulations were conducted by applying a modified version of the generative model (without missing genotypes or errors), using gamete numbers, pruned SNP counts, and crossover counts identical to those inferred from the empirical data. We repeated the above procedure for computing pairwise distances on each simulated dataset, recording the mean proportion of shared alleles across all sperm from each of 25 donors. Repeating this entire procedure 500 times allowed the construction of a null distribution to which we compared our allele sharing calculation from the Sperm-seq data. We then computed a one-tail p-value, defined as the proportion of null simulations where the average pairwise allele sharing was greater than or equal to the observed value.

## Acknowledgments

We thank members of the McCoy lab for feedback and helpful discussions. Thank you to Angela Leung and Alec Wysoker for assistance accessing Sperm-seq data. We also thank the staff at the Maryland Advanced Research Computing Center for computing support.

## Availability of data and materials

Data analysis scripts specific to our study are available at https://github.com/mccoy-lab/transmission-distortion. Our package rhapsodi is available at: https://github.com/mccoy-lab/rhapsodi. Raw sperm sequencing data from Bell et al. [29] can be accessed via dbGaP (study accession number phs001887.v1.p1), as described in the original publication. Raw sperm sequencing data from Leung et al. [30] was accessed upon request from the authors. We filtered the cells in our analysis using metadata published by Bell et al. [29] at: https://zenodo.org/record/3561081#.YLAdO2ZKhb9. Analogous metadata from Leung et al. [30] was obtained upon request from the authors.

## Ethics approval and consent to participate

The Homewood Institutional Review Board of Johns Hopkins University determined that this work does not qualify as federally-regulated human subjects research (HIRB00011435).

## Consent for publication

All authors have read and approved the manuscript.

## Authors’ contributions

S.A.C: Formal Analysis, Funding Acquisition, Investigation, Methodology, Software, Visualization, Writing -Original Draft Preparation, Writing -Review and Editing; K.J.W.: Data Curation, Formal Analysis, Investigation, Methodology, Software, Validation, Visualization, Writing -Original Draft Preparation, Writing -Review and Editing; A.N.B.: Data Curation, Formal Analysis, Investigation, Methodology, Software, Visualization, Writing -Original Draft Preparation, Writing -Review and Editing; D.A.: Formal Analysis, Investigation, Methodology, Writing -Review and Editing; A.D.B.: Data Curation, Methodology, Writing -Review and Editing; R.C.M.: Conceptualization, Funding Acquisition, Methodology, Resources, Software, Supervision, Visualization, Writing -Original Draft Preparation, Writing -Review and Editing

## References

1. Lila Fishman and Mariah McIntosh. Standard deviations: The biological bases of transmission ratio distortion. Annual Review of Genetics, 53(1):347–372, December 2019.

2. Y Koide, Y Shinya, M Ikenaga, N Sawamura, K Matsubara, K Onishi, A Kanazawa, and Y Sano. Complex genetic nature of sex-independent transmission ratio distortion in asian rice species: the involvement of unlinked modifiers and sex-specific mechanisms. Heredity, 108(3):242–247, July 2011.

3. Reflinur, Backki Kim, Sun Mi Jang, Sang-Ho Chu, Yogendra Bordiya, Md Babul Akter, Joohyun Lee, Joong Hyoun Chin, and Hee-Jong Koh. Analysis of segregation distortion and its relationship to hybrid barriers in rice. Rice, 7(1), August 2014.

4. Peng Xu, Jiawu Zhou, Jing Li, Fengyi Hu, Xianneng Deng, Sufeng Feng, Guangyun Ren, Zhi Zhang, Wei Deng, and Dayun Tao. Mapping three new interspecific hybrid sterile loci between oryza sativa and o. glaberrima. Breeding Science, 63(5):476–482, 2014.

5. ZX Tang, XF Wang, MZ Zhang, YH Zhang, DX Deng, and CW Xu. The maternal cytoplasmic environment may be involved in the viability selection of gametes and zygotes. Heredity, 110(4):331–337, November 2012.

6. Yu Yu, Daojun Yuan, Shaoguang Liang, Ximei Li, Xiaqing Wang, Zhongxu Lin, and Xianlong Zhang. Genome structure of cotton revealed by agenome-wide SSR genetic map constructed from a BC1 population between gossypium hirsutum and g. barbadense. BMC Genomics, 12(1), January 2011.

7. Amanda M Hulse-Kemp, Jana Lemm, Joerg Plieske, Hamid Ashrafi, Ramesh Buyyarapu, David D Fang, James Frelichowski, Marc Giband, Steve Hague, Lori L Hinze, Kelli J Kochan, Penny K Riggs, Jodi A Schefier, Joshua A Udall, Mauricio Ulloa, Shirley S Wang, Qian-Hao Zhu, Sumit K Bag, Archana Bhardwaj, John J Burke, Robert L Byers, Michel Claverie, Michael A Gore, David B Harker, Md S Islam, Johnie N Jenkins, Don C Jones, Jean-Marc Lacape, Danny J Llewellyn, Richard G Percy, Alan E Pepper,\ Jesse A Poland, Krishan Mohan Rai, Samir V Sawant, Sunil Kumar Singh, Andrew Spriggs, Jen M Taylor, Fei Wang, Scott M Yourstone, Xiuting Zheng, Cindy T Lawley, Martin W Ganal, Allen Van Deynze, Iain W Wilson, and David M Stelly. Development of a 63k SNP array for cotton and high-density mapping of intraspecific and interspecific populations of gossypium spp. G3 Genes Genomes Genetics, 5(6):1187–1209, June 2015.

8. Baosheng Dai, Huanle Guo, Cong Huang, Muhammad M. Ahmed, and Zhongxu Lin. Identification and characterization of segregation distortion loci on cotton chromosome 18. Frontiers in Plant Science, 7, January 2017.

9. Amanda M Larracuente and Daven C Presgraves. The selfish segregation distorter gene complex of drosophila melanogaster. Genetics, 192(1):33–53, September 2012.

10. S. R. Mcdermott and M. A. F. Noor. Mapping of within-species segregation distortion in drosophila persimilis and hybrid sterility between d. persimilis and d. pseudoobscura. Journal of Evolutionary Biology, 25(10):2023–2032, August 2012.

11. Josephine A. Reinhardt, Cara L. Brand, Kimberly A. Paczolt, Philip M. Johns, Richard H. Baker, and Gerald S. Wilkinson. Meiotic drive impacts expression and evolution of x-linked genes in stalk-eyed flies. PLoS Genetics, 10(5):e1004362, May 2014.

12. Chevonne D Eversley, Tavia Clark, Yuying Xie, Jill Steigerwalt, Timothy A Bell, Fernando PM de Villena, and David W Threadgill. Genetic mapping and developmental timing of transmission ratio distortion in a mouse interspecific backcross. BMC Genetics, 11(1):98, 2010.

13. Joaquim Casellas, Rodrigo J Gularte, Charles R Farber, Luis Varona, Margarete Mehrabian, Eric E Schadt, Aldon J Lusis, Alan D Attie, Brian S Yandell, and Juan F Medrano. Genome scans for transmission ratio distortion regions in mice. Genetics, 191(1):247–259, May 2012.

14. Lisa E Kursel and Harmit S Malik. The cellular mechanisms and consequences of centromere drive. Current Opinion in Cell Biology, 52:58–65, June 2018.

15. María Angélica Bravo Núńez, Ibrahim M. Sabbarini, Michael T. Eickbush, Yue Liang, Je↵rey J. Lange, Aubrey M. Kent, and Sarah E. Zanders. Dramatically diverse schizosaccharomyces pombe wtf meiotic drivers all display high gamete-killing effciency. PLOS Genetics, 16(2):e1008350, February 2020.

16. David Bikard, Dhaval Patel, Claire L. Metté, Veronica Giorgi, Christine Camilleri, Malcolm J Bennett, and Olivier Loudet. Divergent evolution of duplicate genes leads to genetic incompatibilities within a. thaliana. Science, 323(5914):623–626, 2009.

17. Kenneth G. Ross and DeWayne Shoemaker. Unexpected patterns of segregation distortion at a selfish supergene in the fire ant solenopsis invicta. BMC Genetics, 19(1), November 2018.

18. John Schimenti. Segregation distortion of mouse t haplotypes. Trends in Genetics, 16(6):240–243, June 2000.

19. David M Higgins, Elizabeth G Lowry, Lisa B Kanizay, Philip W Becraft, David W Hall, and R Kelly Dawe. Fitness costs and variation in transmission distortion associated with the abnormal chromosome 10 meiotic drive system in maize. Genetics, 208(1):297–305, January 2018.

20. N. Phadnis and H. A. Orr. A single gene causes both male sterility and segregation distortion in drosophila hybrids. Science, 323(5912):376–379, January 2009.

21. Adele A Mitchell, David J Cutler, and Aravinda Chakravarti. Undetected genotyping errors cause apparent overtransmission of common alleles in the transmission/disequilibrium test. The American Journal of Human Genetics, 72(3):598–610, 2003.

22. Sebastian Zöllner, Xiaoquan Wen, Neil A. Hanchard, Mark A. Herbert, Carole Ober, and Jonathan K. Pritchard. Evidence for extensive transmission distortion in the human genome. The American Journal of Human Genetics, 74(1):62–72, January 2004.

23. Wynn K Meyer, Barbara Arbeithuber, Carole Ober, Thomas Ebner, Irene Tiemann-Boege, Richard R Hudson, and Molly Przeworski. Evaluating the evidence for transmission distortion in human pedigrees. Genetics, 191(1):215–232, May 2012.

24. Richard S Spielman, Ralph E McGinnis, and Warren J Ewens. Transmission test for linkage disequilibrium: the insulin gene region and insulin-dependent diabetes mellitus (iddm). American journal of human genetics, 52(3):506, 1993.

25. Andrew D Paterson, Daryl Waggott, Arne Schillert, Claire Infante-Rivard, Shelley B Bull, Yun Joo Yoo, and Dushanthi Pinnaduwage. Transmission-ratio distortion in the framingham heart study. In BMC proceedings, volume 3, issue 7, pages 1–5. BioMed Central, 2009.

26. Jianbin Wang, H Christina Fan, Barry Behr, and Stephen R Quake. Genome-wide single-cell analysis of recombination activity and de novo mutation rates in human sperm. Cell, 150(2):402–412, 2012.

27. Linda Odenthal-Hesse, Ingrid L Berg, Amelia Veselis, Alec J Je↵reys, and Celia A May. Transmission distortion a↵ecting human noncrossover but not crossover recombination: a hidden source of meiotic drive. PLoS genetics, 10(2):e1004106, 2014.

28. Graham Coop and Simon R Myers. Live hot, die young: transmission distortion in recombination hotspots. PLoS genetics, 3(3):e35, 2007.

29. Avery Davis Bell, Curtis J. Mello, James Nemesh, Sara A. Brumbaugh, Alec Wysoker, and Steven A. McCarroll. Insights into variation in meiosis from 31, 228 human sperm genomes. Nature, 583(7815):259–264, June 2020.

30. Angela Q Leung, Avery Davis Bell, Curtis J Mello, Alan S Penzias, Steven A McCarroll, and Denny Sakkas. Single cell analysis of dna in more than 10,000 individual sperm from men with abnormal reproductive outcomes. Journal of Assisted Reproduction and Genetics, pages 1–9, 2021.

31. The International SNP Map Working Group. A map of human genome sequence variation containing 1.42 million single nucleotide polymorphisms. Nature, 409:928–933, 2001.

32. Karl W Broman and James L Weber. Characterization of human crossover interference. The American Journal of Human Genetics, 66(6):1911–1926, 2000.

33. Ruidong Li, Han Qu, Jinfeng Chen, Shibo Wang, John M Chater, Le Zhang, Julong Wei, Yuan-Ming Zhang, Chenwu Xu, Wei-De Zhong, Jianguo Zhu, Jianming Lu, Yuanfa Feng, Weiming Chen, Renyuan Ma, Sergio Pietro Ferrante, Mikeal L Roose, and Zhenyu Jia. Inference of chromosome-length haplotypes using genomic data of three or a few more single gametes. Molecular Biology and Evolution, 37(12):3684–3698, July 2020.

34. 1000 Genomes Project Consortium et al. A global reference for human genetic variation. Nature, 526(7571):68, 2015.

35. Karl W Broman, Je↵rey C Murray, Val C Sheffeld, Raymond L White, and James L Weber. Comprehensive human genetic maps: individual and sex-specific variation in recombination. The American Journal of Human Genetics, 63(3):861–869, 1998.

36. Xiaoyi Gao, Joshua Starmer, and Eden R Martin. A multiple testing correction method for genetic association studies using correlated single nucleotide polymorphisms. Genetic Epidemiology: The Oicial Publication of the International Genetic Epidemiology Society, 32(4):361–369, 2008.

37. Xiaoyi Gao, Lewis C Becker, Diane M Becker, Joshua D Starmer, and Michael A Province. Avoiding the high bonferroni penalty in genome-wide association studies. Genetic Epidemiology: The Oicial Publication of the International Genetic Epidemiology Society, 34(1):100–105, 2010.

38. Joseph H Marcus and John Novembre. Visualizing the geography of genetic variants. Bioinformatics, 33(4):594–595, 2017.

39. Frances E Clark and Takashi Akera. Unravelling the mystery of female meiotic drive: where we are. Open Biology, 11(9):210074, 2021.

40. Joe H Ward Jr. Hierarchical grouping to optimize an objective function. Journal of the American statistical association, 58(301):236–244, 1963.

41. G David Forney. The viterbi algorithm. Proceedings of the IEEE, 61(3):268–278, 1973.

42. Yu Hou, Wei Fan, Liying Yan, Rong Li, Ying Lian, Jin Huang, Jinsen Li, Liya Xu, Fuchou Tang, X. Sunney Xie, and Jie Qiao. Genome analyses of single human oocytes. Cell, 155(7):1492–1506, December 2013.

43. Marcel Martin, Murray Patterson, Shilpa Garg, Sarah Fischer, Nadia Pisanti, Gunnar W Klau, Alexander Schöenhuth, and Tobias Marschall. Whatshap: fast and accurate read-based phasing. BioRxiv, page 085050, 2016.

44. Peter Edge, Vineet Bafna, and Vikas Bansal. Hapcut2: robust and accurate haplotype assembly for diverse sequencing technologies. Genome research, 27(5):801–812, 2017.

45. Peter Krusche,, Len Trigg, Paul C. Boutros, Christopher E. Mason, Francisco M. De La Vega, Benjamin L. Moore, Mar Gonzalez-Porta, Michael A. Eberle, Zivana Tezak, Samir Lababidi, Rebecca Truty, George Asimenos, Birgit Funke, Mark Fleharty, Brad A. Chapman, Marc Salit, and Justin M. Zook. Best practices for benchmarking germline small-variant calls in human genomes. Nature Biotechnology, 37(5):555–560, March 2019.

46. Haley M. Amemiya, Anshul Kundaje, and Alan P. Boyle. The ENCODE blacklist: Identification of problematic regions of the genome. Scientific Reports, 9(1), June 2019.

47. Shaun Purcell, Benjamin Neale, Kathe Todd-Brown, Lori Thomas, Manuel AR Ferreira, David Bender, Julian Maller, Pamela Sklar, Paul IW De Bakker, Mark J Daly, et al. Plink: a tool set for whole-genome association and population-based linkage analyses. The American journal of human genetics, 81(3):559–575, 2007.

48. David Porubsky, Shilpa Garg, Ashley D. Sanders, Jan O. Korbel, Victor Guryev, Peter M. Lansdorp, and Tobias Marschall. Dense and accurate whole-chromosome haplotyping of individual genomes. Nature Communications, 8(1), November 2017.

