## Supplementary Materials for "Strict adherence to Mendel’s First Law across a large sample of human sperm genomes"

### Definition of metrics

- Accuracy (Phasing & Imputation): For two sequences of genotypes, s1 and s2, (of equal length and both represented by 0s, 1s, and NAs), accuracy was defined as the Hamming error rate subtracted from 1. The Hamming error rate was defined to be equal to the number of positions with mismatches between s1 and s2 (ignoring a mismatch when a single sequence had an NA value) divided by the total number of sequence positions [33, 48].
- Completeness: For a sequence of genotypes, s1 (represented by 0s, 1s, and NAs), completeness was defined as the number of non-NA positions divided by the total number of positions in s1 [33].
- Switch Error Rate: For two sequences of genotypes, s1 and s2 (of equal length and both represented by 0s, 1s, and NAs), the switch error rate was defined as the number of first mismatches in any stretch of adjacent mismatches (where the stretch is greater than or equal to length 1 and uninterrupted by NAs) divided by the total number of sequence positions [33].
- Largest Haplotype Segment: For two sequences of genotypes, s1 and s2 (of equal length and both represented by 0s, 1s, and NAs), the largest haplotype segment was defined as the maximum number of adjacent matched positions, uninterrupted by NAs. For plotting purposes, we divide this number by the total number of sequence positions [33].
- True Positive (TP): A true simulated meiotic recombination breakpoint intersects with predicted meiotic recombination breakpoint (Fig. S3b).
- True Negative (TN): Simulated chromosome with no meiotic recombination events is correctly predicted to have no recombination breakpoints (Fig. S3b).
- False Positive (FP): A predicted meiotic recombination breakpoint does not intersect any true simulated meiotic recombination breakpoints (Fig. S3b).
- False Negative (FN): A true simulated meiotic recombination breakpoint does not intersect any predicted meiotic recombination breakpoints (Fig. S3b).
- Recall or True Positive Rate (TPR): $\frac{TP}{TP+FN}$
- Precision: $\frac{TP}{TP+FP}$
- F1 Score: $\frac{2 \times precision \times recall}{precision + recall}$
- False Discovery Rate (FDR): $\frac{FP}{TP+FP}$
- False Positive Rate (FPR): $\frac{FP}{TN+FP}$
- Specificity or True Negative Rate (TNR): $\frac{TN}{TN+FP}$
- Accuracy (Discovery): $\frac{TP+TN}{TP+TN+FP+FN}$

### Power analysis for detecting TD

To evaluate the statistical power of our TD scanning approach, we conducted simulations of various levels of TD (0-10% deviations from Mendelian expectations) across a range of sample sizes of human sperm (Fig. S13). Applying binomial tests to these simulated data, we find that in a sample of more than 1000 sperm, our approach has high statistical power (>80%) to detect even subtle transmission distortion at a single locus. Larger samples increase our power to detect even smaller deviations from binomial expectations. The power was computed for each study design (number of gametes and rate of transmission distortion) from 1000 independent trials.

### Genotype filtering to mitigate spurious TD signatures

We removed any sperm cells designated as poor quality and any cell that was called as aneuploid for the chromosome of interest by the original researchers. We conduct this filtering based on metadata from [29] (published) and [30] (obtained on request).

To limit potential artifacts, we excluded all technically challenging regions of the genome previously identified by the Genome in a Bottle Consortium (GRCh38 "union") [45] and/or the ENCODE Consortium [46]. These include tandem repeats, homopolymers >6bp, imperfect homopolymers >10bp, difficult to map regions, segmental duplications, GC content <25% or >65%, and other difficult regions.

As further filtering at the level of individual variants, we restricted our analysis to SNPs that are also present in the 1,000 Genomes Project dataset [34], which should be enriched for genuine variation in comparison to singleton variation that is private to the donor individuals. Notably, even if TD were to be caused by a rare variant (which we would have removed from our dataset in this step), it would likely occur on a haplotype that also carries common variants, allowing us to discern its effect.

Finally, we remove any SNPs exhibiting an excess of observations across sperm cells (>1 standard deviation above the mean), with the goal of filtering out potential segmental duplications that are private to a given donor. Such duplications contribute to alignment artifacts, whereby both copies of the sequence (including divergent sites) would pile up at the same location. Such phenomena may generate false signatures of TD, which we detail here.

Specifically, we identified two potential indicators that a given signature of TD may arise from rare segmental duplications (Fig. S14). First, we consider the number of sperm in which the SNP is observed to carry the reference allele vs. the alternative allele (Fig. S14a, Fig. S14b). In the absence of TD, we expect points (each representing a SNP) to cluster to the line with slope = 1, showing even representation of both the reference and the alternative allele for that SNP in a donor's pool of sperm. Such is the case in Fig. S14b, which is representative of most loci throughout the genome. However, if no filtering is applied, we observe that certain regions with strong TD signatures exhibit predictable patterns of allelic imbalance (manifesting here as lines with slope = 0.5 and slope = 2 (Fig. S14a).

The second indicator that such regions derive from a rare segmental duplication regards the overall number of genotype observations. Because of the low coverage ($\sim 0.01\times$), we expect most SNPs to be present in only a few of the 969-3,377 sperm per donor. If no filtering is applied, we find that SNPs exhibiting strong signatures of TD also exhibit an extreme excess of genotype observations, again consistent with duplication.

To avoid these potential confounding impacts of segmental duplications, we set a stringent threshold for our filtering pipeline, removing SNPs with excess (>1 standard deviation above the mean) genotype observations across the pool of sperm. We calculated these means and standard deviations separately for each donor's chromosomes (25 donors $\times$ 22 chromosomes).

### Simulation of TD

To simulate TD signatures, we developed a modified version of our generative model, whereby we randomly select a SNP and assign one of the haplotypes as a deleterious allele (Fig. S11). We then generate a larger than desired set of gametes, remove a user-specified fraction of gametes containing this SNP/haplotype combination from the dataset, and sample the desired number of gametes from this pool.

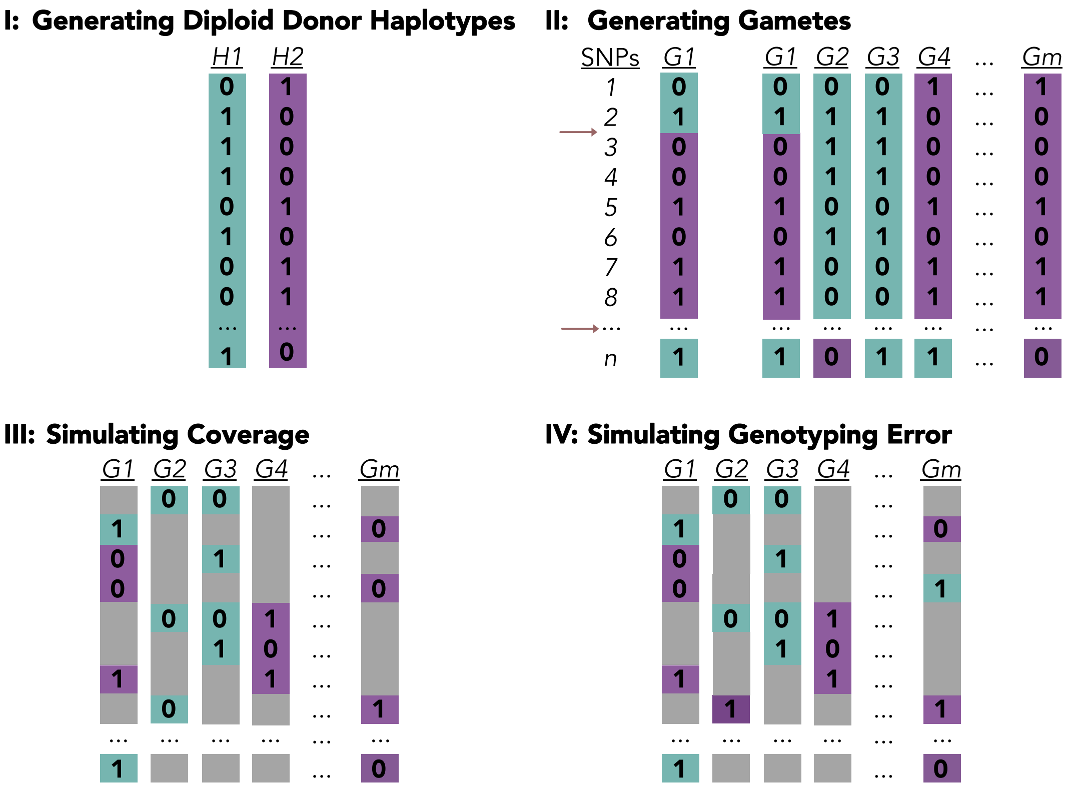


Figure S1 Generative Model. (I) The first step of the generative model builds the phased haplotypes of the diploid donor, with n hetSNPs. (II) In the second step, gamete genotypes are derived from the diploid donor using a no-interference model for each gamete chromosome, repeating the process for all gametes, {1, ..., m}. (III) In the third step, low-coverage sequencing data is generated by removing genotypes from a copy of the gamete genotype matrix. (IV) In the final step, genotyping error is simulated by replacing correct genotypes with the opposite allele.

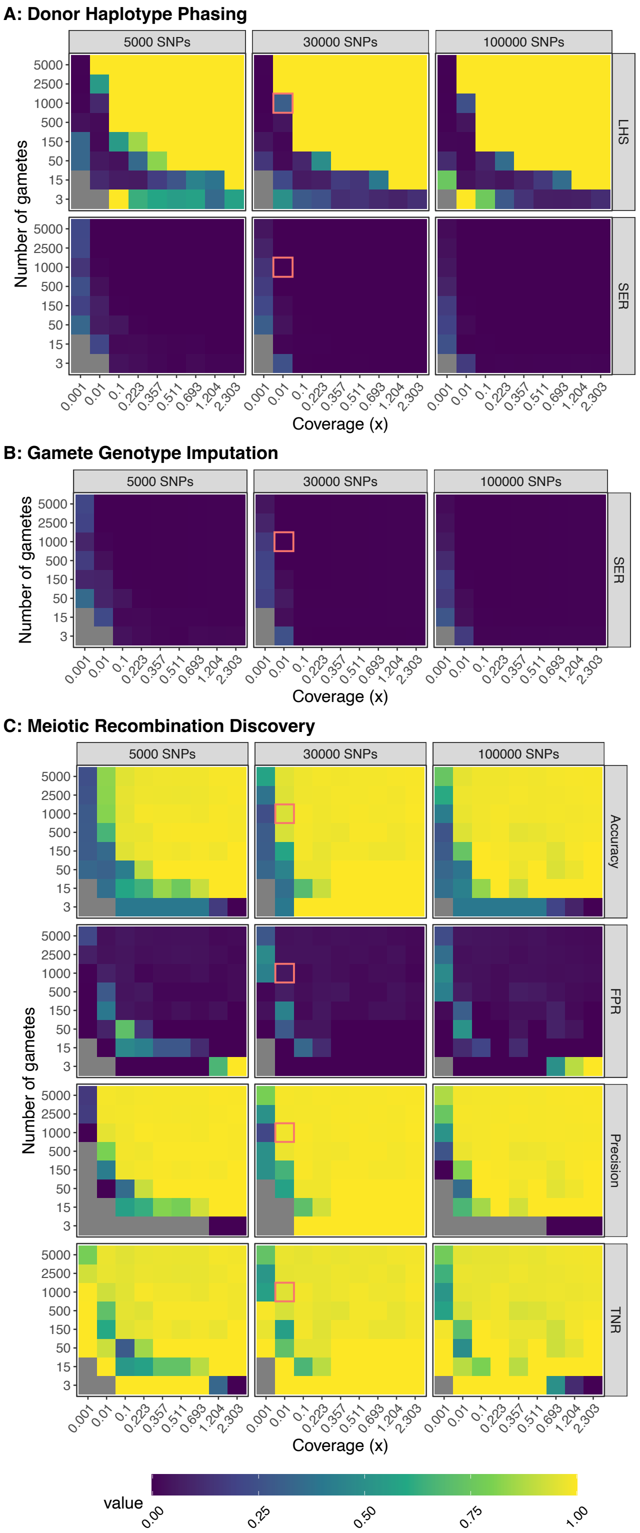


Figure S2 Benchmarking performance across a wide range of input data profiles - Additional Metrics. Input data were created from the generative model and analyzed with rhapsodi. For all data in this figure, the genotyping error and recombination rates were matched between models. Each value is the average of three independent trials. LHS: Largest Haplotype Segment (as ratio of segment length / total hetSNPs); SER: Switch Error Rate; FPR: False Positive Rate; TNR: True Negative Rate.

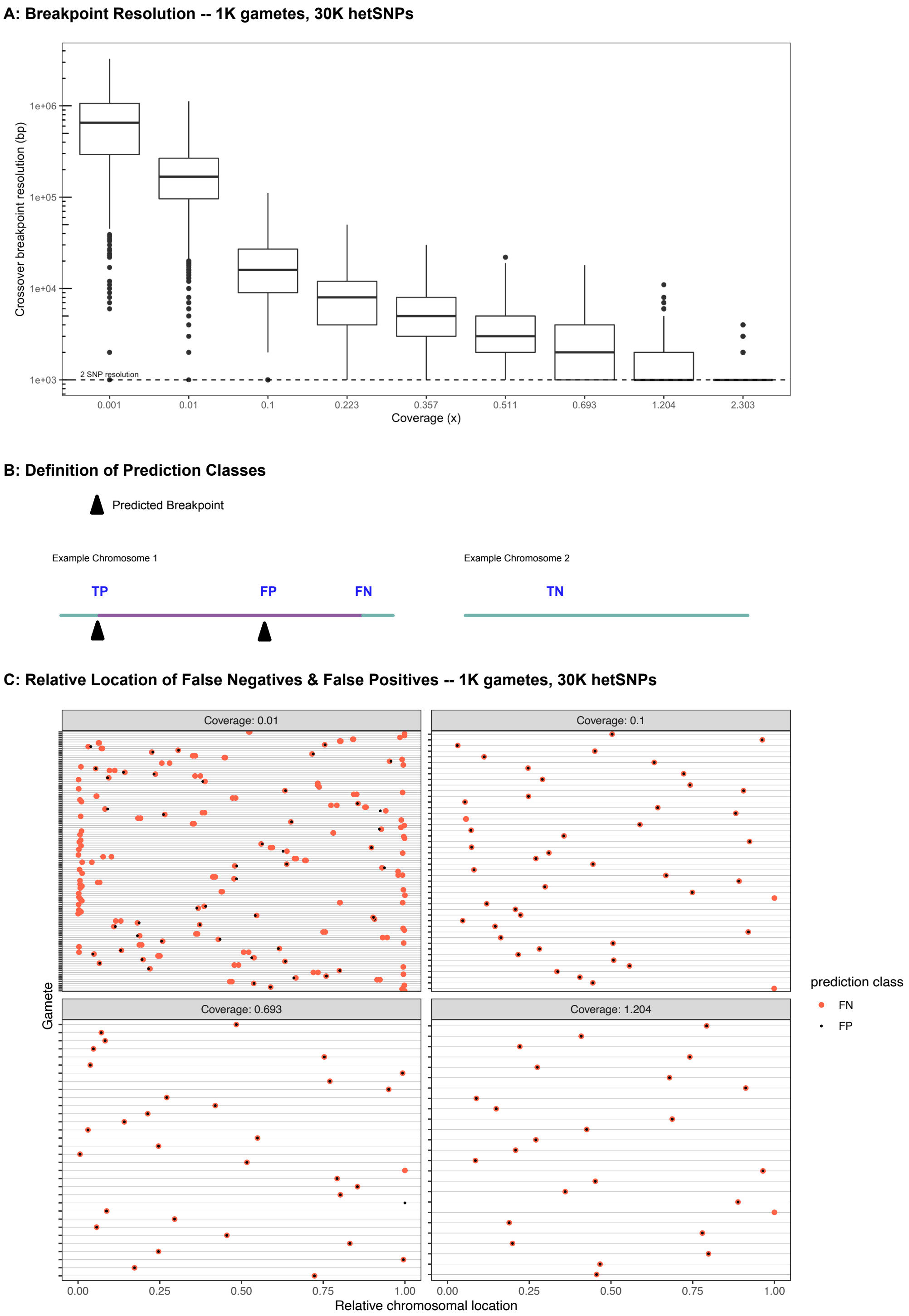


Figure S3 Discovery of meiotic recombination events in simulated data profiles reflecting Sperm-seq data. (A) Breakpoint resolution. Pooling all rhapsodi predicted breakpoints, data is stratified by sequencing coverage. Breakpoint resolution in bp is approximated by 1000 × (the number of hetSNPs in the breakpoint − 1). Because "ground truth" breakpoints are 2 adjacent hetSNPs, the dashed-line is y=1000. (B) Definition of prediction classes. True positive (TP); True negative (TN); False positive (FP); & False negative (FN). Chromosome color represents inherited donor haplotype; change in color depicts a recombination event. (C) Relative location of FN & FP breakpoints within gametes from selected, simulated coverages. Each row in each panel represents an individual gamete with at least one FN or FP. Relative SNP location is index / total hetSNPs.

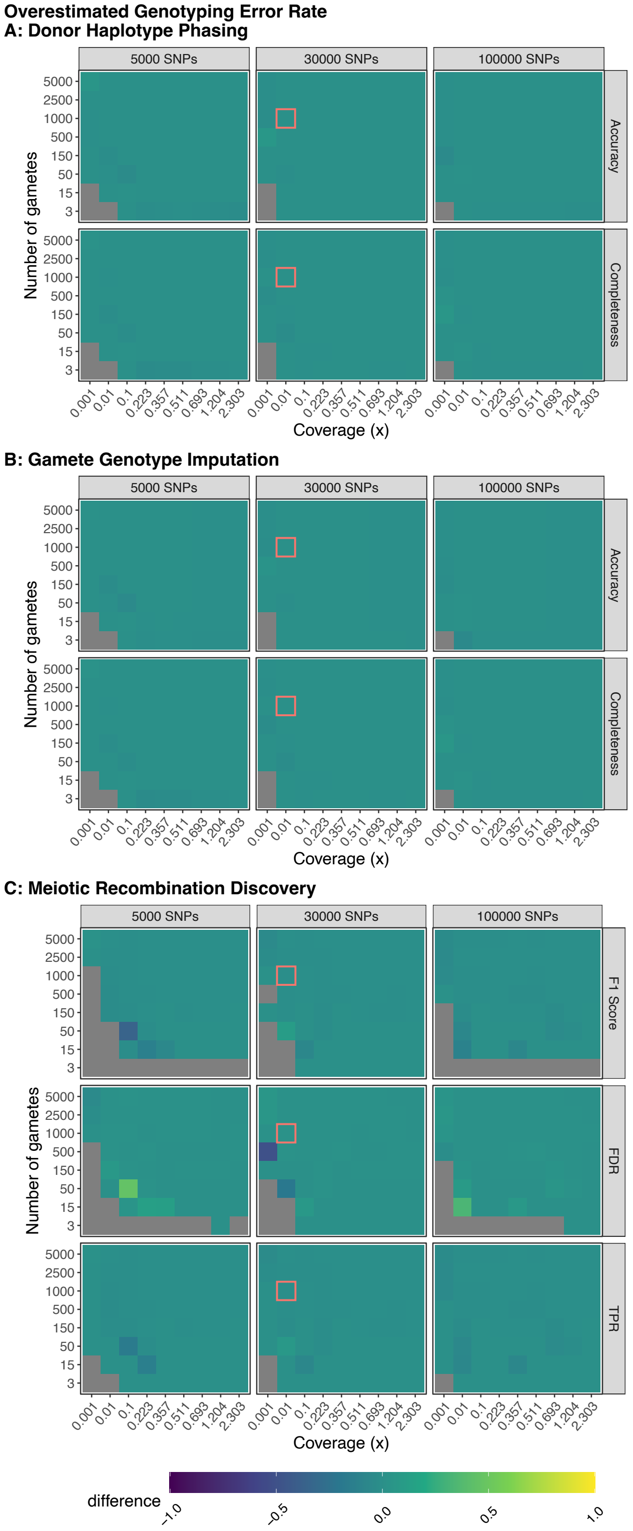


Figure S4 Model robustness when genotyping error is overestimated. Plotted values are the difference between the mean performance (across 3 independent trials) when the generative model and rhapsodi use the same parameters (Fig. 2) and the mean performance (across 3 independent trials) when the rhapsodi genotyping error rate (0.005) is overestimated compared to the rate used by the generative model (0.001). The recombination rate is 1 in both models.

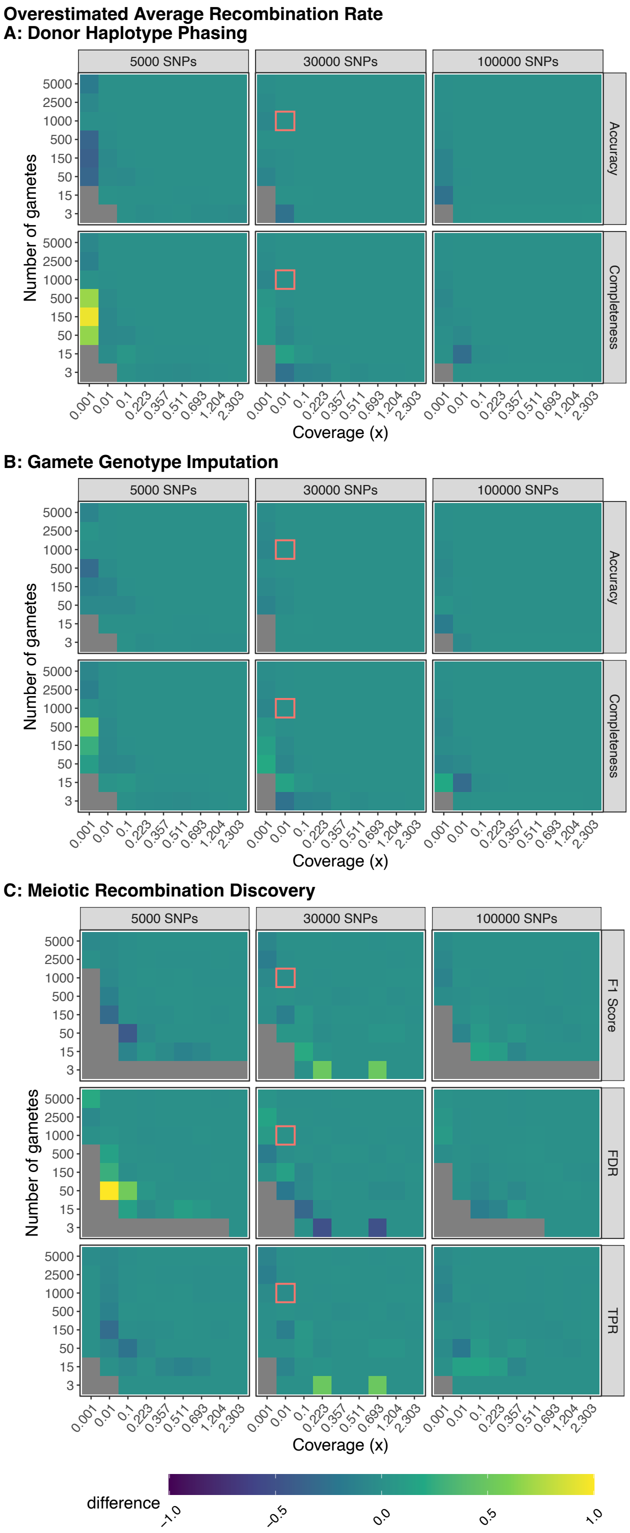


Figure S5 Model robustness when recombination rate is overestimated. Plotted values are the difference between the mean performance (across 3 independent trials) when the generative model and rhapsodi use the same parameters (Fig. 2) and the mean performance (across 3 independent trials) when the rhapsodi recombination rate (1) is overestimated compared to the rate used by the generative model (0.6). The error rate is 0.005 in both models.

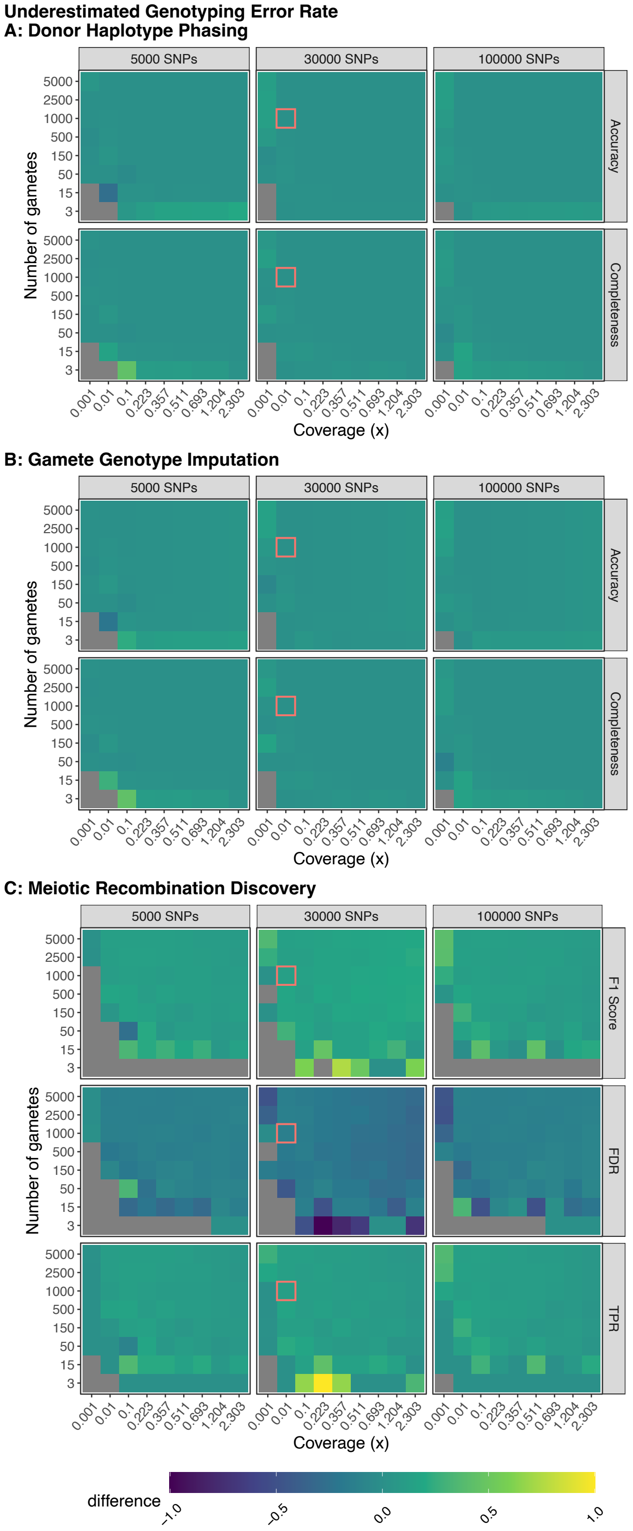


Figure S6 Model robustness when genotyping error is underestimated. Plotted values are the difference between the mean performance (across 3 independent trials) when the generative model and rhapsodi use the same parameters (Fig. 2) and the mean performance (across 3 independent trials) when the rhapsodi genotyping error (0.005) is underestimated compared to the rate used by the generative model (0.05). The recombination rate is 1 in both models.

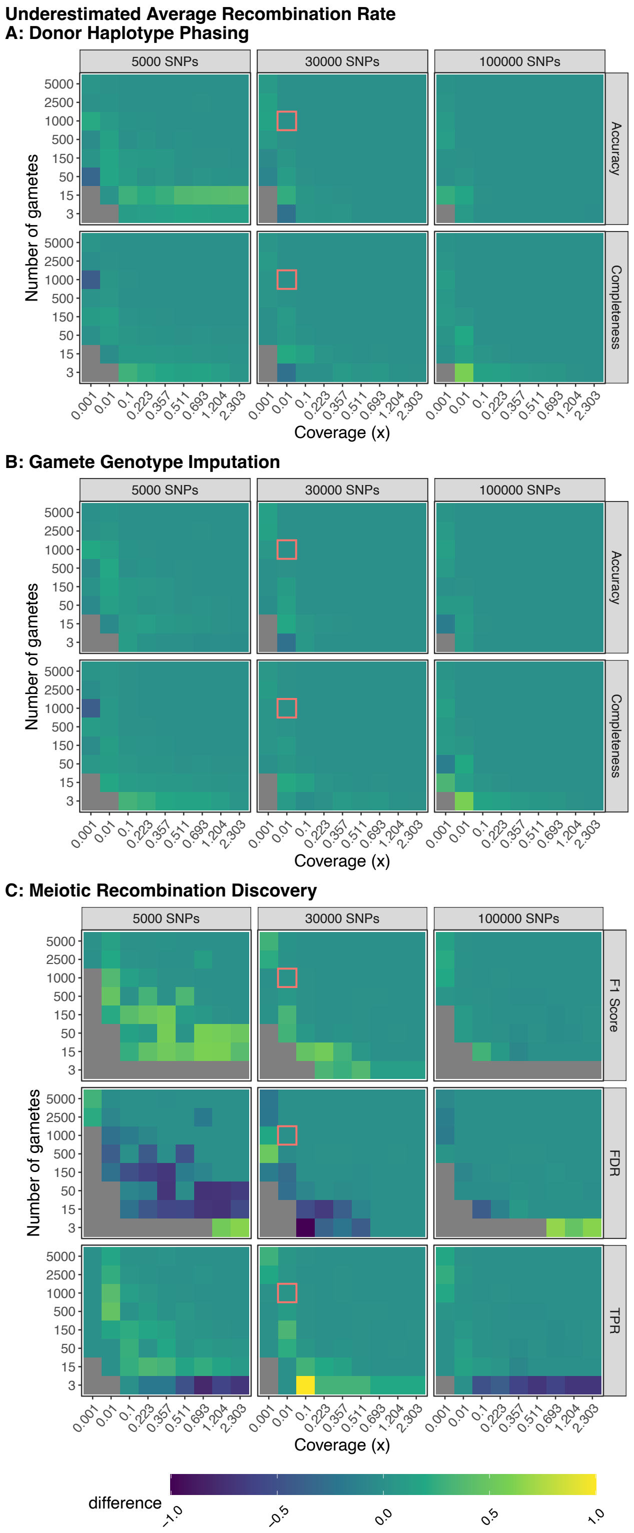


Figure S7 Model robustness when recombination rate is underestimated. Plotted values are the difference between the mean performance (across 3 independent trials) when the generative model and rhapsodi use the same parameters (Fig. 2) and the mean performance (across 3 independent trials) when the rhapsodi recombination rate (1) is underestimated compared to the rate used by the generative model (3). The error rate is 0.005 in both models.

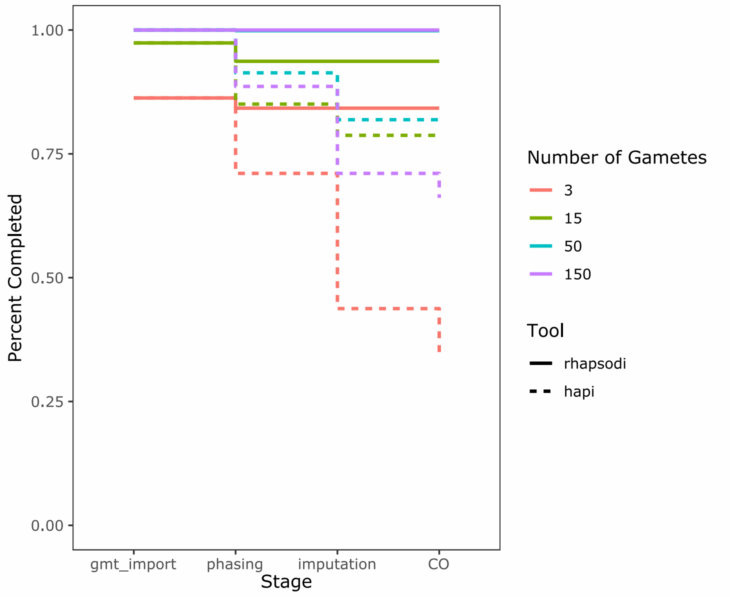


Figure S8 Comparison of the percentage of simulated datasets successfully analyzed by rhapsodi and Hapi across (1) Data import (gmt_import) (2) Donor haplotype phasing (phasing) (3) Gamete genotype imputation (imputation), & (4) Meiotic recombination discovery (CO) stages. The datasets were stratified into groups according to the number of gametes used in construction, signified by color. The run completion trend for each tool is signified by line-type.

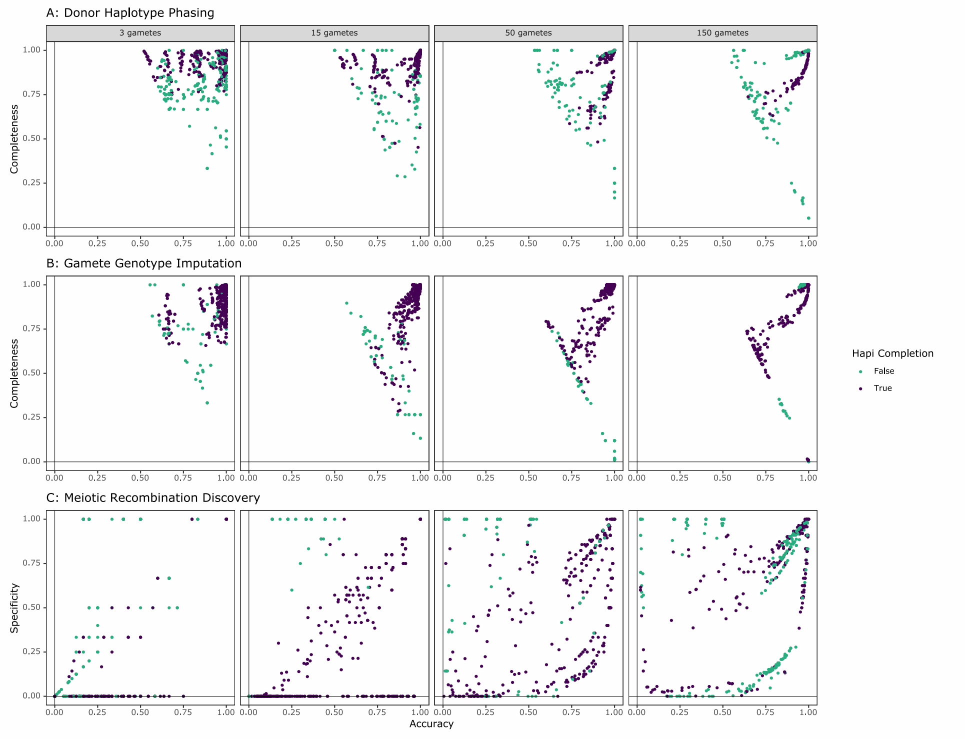


Figure S9 Performance of rhapsodi on each simulated dataset, colored based on Hapi's ability to successfully analyze each given data set. False = Not successful; True = Successful. Each column is defined by the number of gametes used in data construction. (A) The association between phasing accuracy and completeness. (B) The association between mean gamete imputation accuracy and mean completeness. (C) The association between meiotic recombination discovery accuracy and specificity.

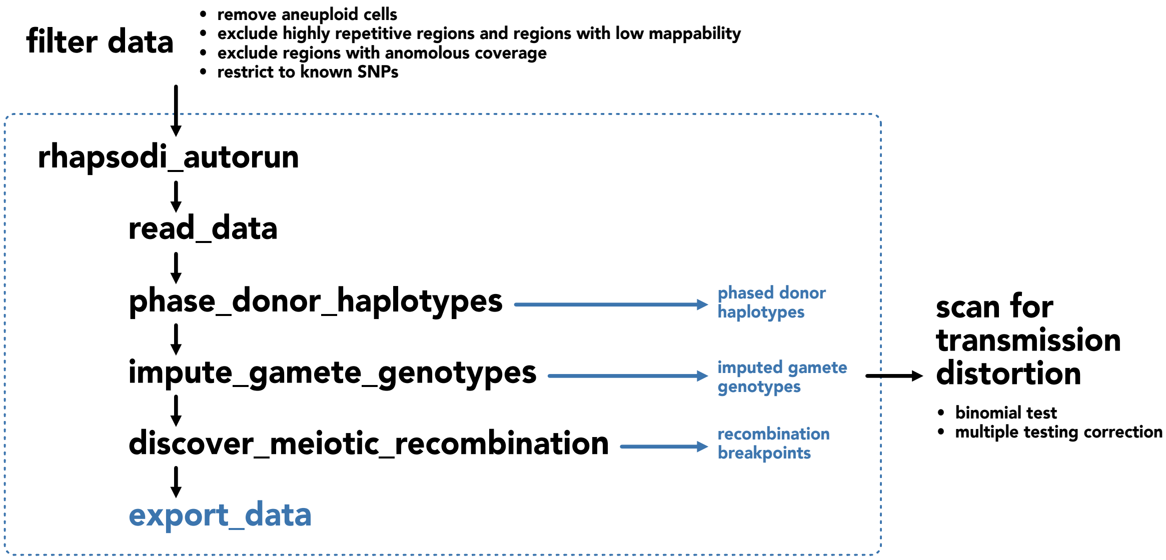


Figure S10 Workflow for application of rhapsodi and investigation of transmission distortion. Raw data containing hetSNPs from each chromosome are filtered to limit spurious signal caused by sequencing error. The autorun function from rhapsodi is applied, which generates and exports data from phased donor haplotypes, imputed gamete genotypes, and meiotic recombination breakpoints. Using the imputed gamete genotypes (generated with the unsmoothed option in rhapsodi), we conduct a binomial test for transmission distortion. We compare our results with our genome-wide threshold for statistical significance, which takes into account multiple hypothesis testing and the extreme linkage disequilibrium in these data.

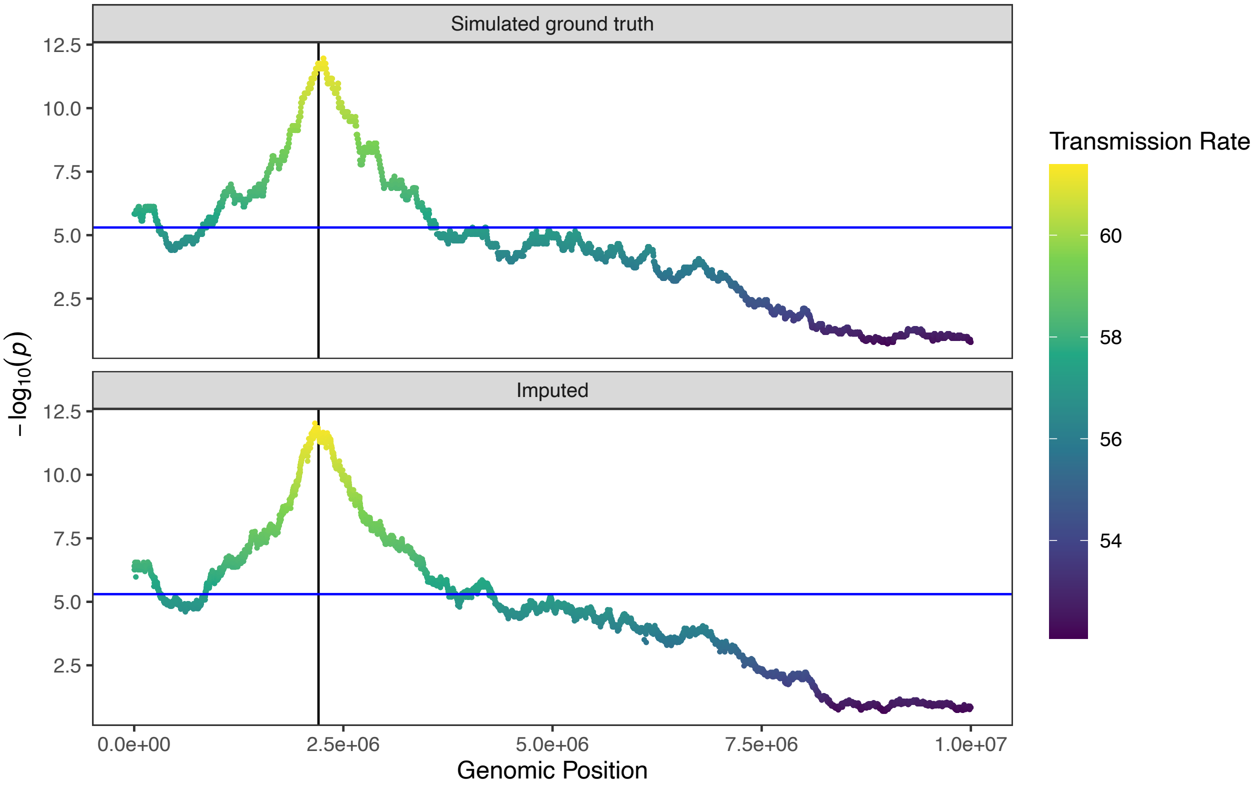


Figure S11 Simulated signature of transmission distortion. We simulated 1,000 gametes with 10,000 SNPs. We generated TD by choosing an allele at random and removing 30% of gametes which carried that allele. We simulated coverage of 0.01× and genotyping error rate of 0.005 and then imputed gamete genotypes using rhapsodi. Top and bottom panels shows results of testing for transmission distortion on simulated ground truth and imputed data, respectively.

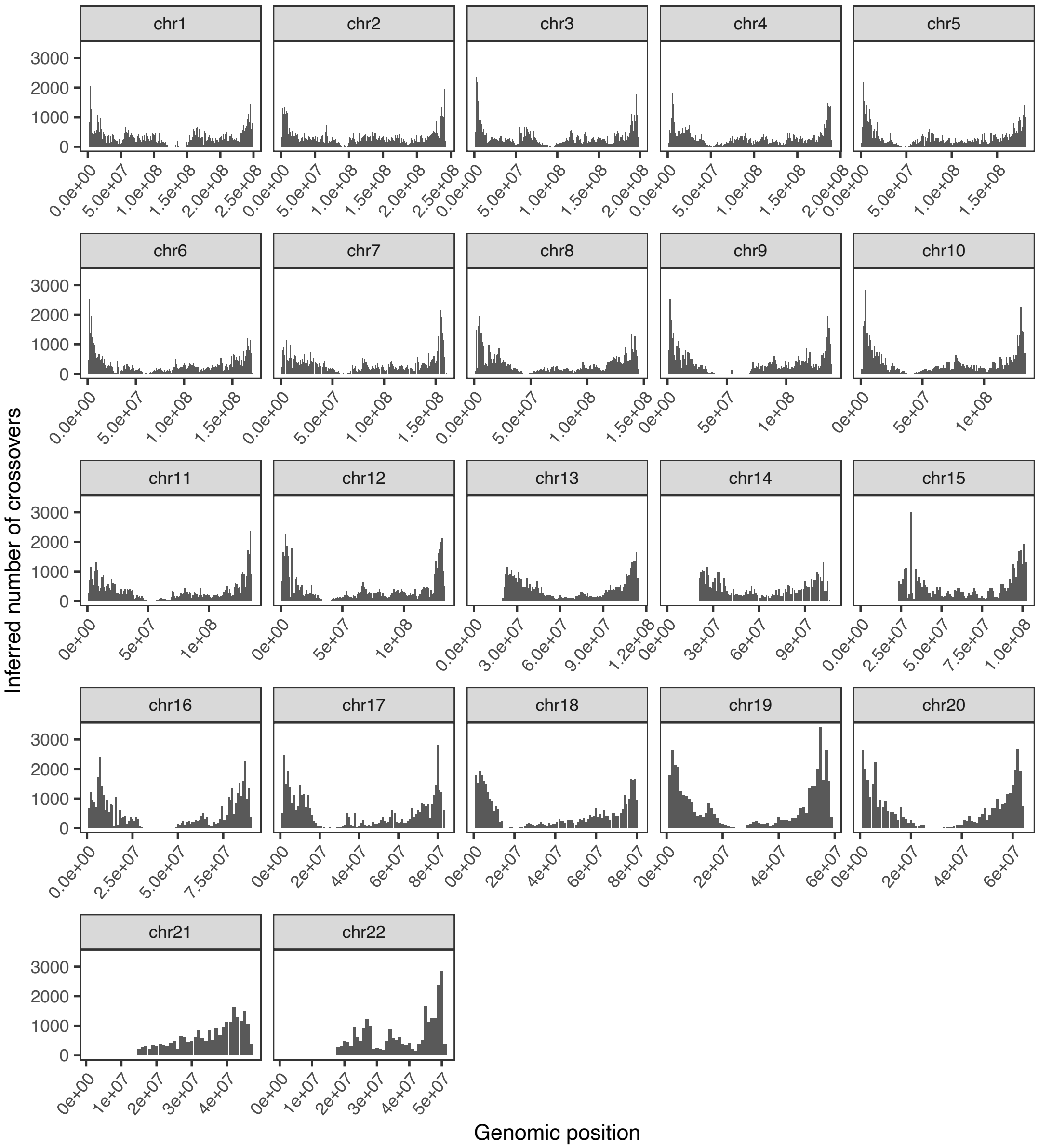


Figure S12 Recombination map of inferred crossovers in the Sperm-seq data. This map displays counts of the inferred number of crossovers within each 1 Mbp bin for each chromosome, pooling inference across the 25 donors. We used the midpoint of each crossover's predicted breakpoints to localize the event to a specific bin.

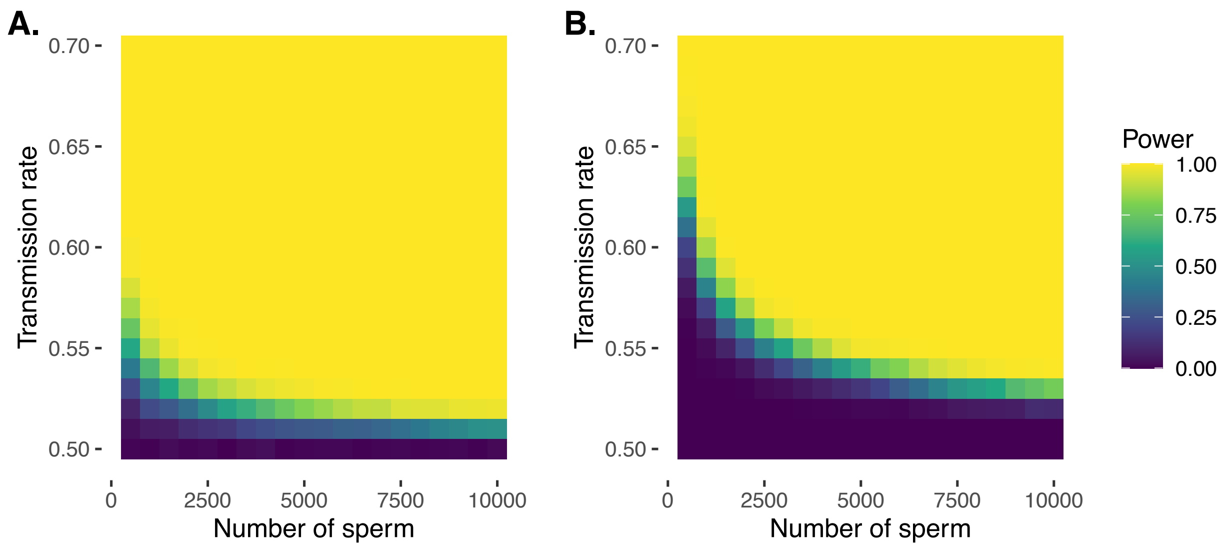


Figure S13 Simulation demonstrating power to detect deviations from binomial expectations across sample sizes of sperm, without (A) and with (B) multiple testing correction. The power for each study design was computed from 1000 independent simulations. For panel B, the Bonferroni-adjusted p-value threshold of $1.78 \times 10^{-7}$ was used for consistency with data analysis.

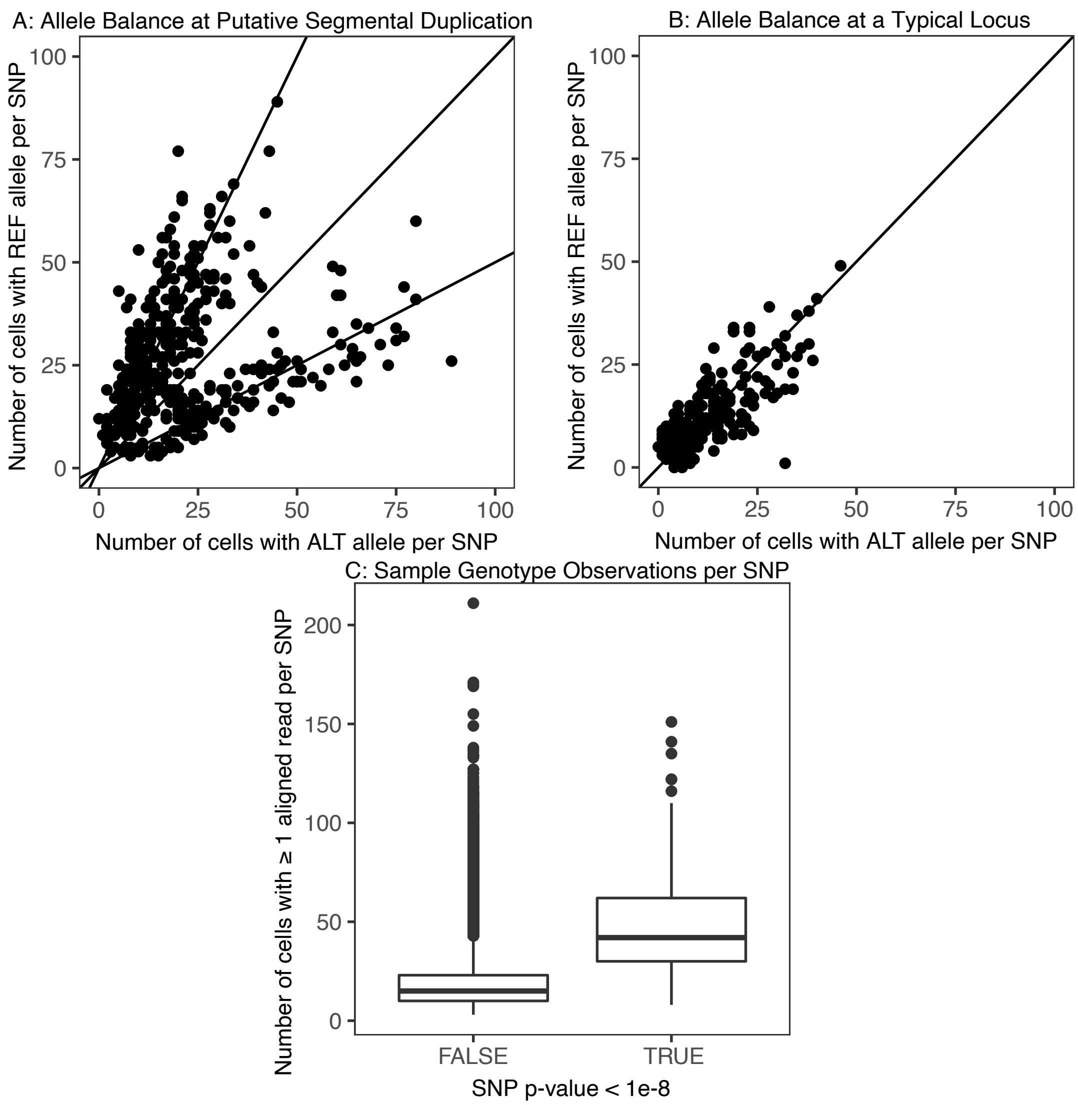


Figure S14 Example of evidence for segmental duplication in donor NC17, chromosome 6. (A) For a segment of the chromosome enriched for low uncorrected p-values (p-value $< 1 \times 10^{-8}$), we plot each hetSNP based on the number of sperm in which it was observed as the reference allele or alternative allele. Points cluster on lines with slope = 0.5 and slope = 2, deviating from the null expectation of clustering on slope = 1 (equal representation of the alternative and reference alleles across the pool of sperm). (B) To compare, we consider SNPs across the rest of the chromosome, outside of this region. Points cluster to the line slope = 1. (C) Across the whole chromosome, we consider the number of sperm cells in which each SNP is observed, based on whether their uncorrected p-value was greater or less than $1 \times 10^{-8}$.
